# Unbiased and scalable reduction of diverse bacterial genomes

**DOI:** 10.64898/2026.09.02.748983

**Authors:** Mikel Lipschitz, Baiyi Quan, Indeever Madireddy, Gohta Aihara, Marisa Bennett, Tsui-Fen Chou, Kaihang Wang

## Abstract

The genome is a complex, integrated system where the functions and regulatory interactions of its many components remain poorly understood. Genome minimization aims to reduce genomic complexity by removing non-essential elements to reveal the fundamental building blocks of cellular life. However, current minimization strategies are often slow and species-specific due to a reliance on prior information, and limited to producing single, isolated strains, which obscures the diverse ways a genome can adapt to large-scale DNA removal. Here we show the development and application of Stochastic Lineage-based Iterative Minimization (SLIM) a modular, high-throughput platform for unbiased genome reduction across phylogenetically diverse bacteria. We apply SLIM to generate a library of genome-reduced *Escherichia coli* lineages. We then interrogate the lineages, identifying both universal and lineage-specific transcriptional and translational reprogramming in response to deletions. We demonstrate that these expression dynamics drive environment-dependent fitness, allowing us to pinpoint a single gene deletion in one genome-reduced lineage as the driver of a measurable environmental growth defect. Beyond *E. coli*, we successfully deploy SLIM in phylogenetically distinct bacterial taxa to rapidly reduce the genomes of *Shigella flexneri* and *Pseudomonas putida*, distinct genus and order respectively from *E. coli*, without species-specific optimization. Our results establish a scalable, generalizable framework for navigating the vast landscape of minimized genomes, providing a powerful new tool for functional discovery and the rational design of synthetic genomic chassis.

## Main

The genome is a paradox; a nearly invisible system so robust it orchestrates every trait of every living thing, yet so fragile that a single change can spell certain death. Genes, the protein-coding elements of the genome, encode a variety of functions and features. Regulatory and other non-coding regions determine when, where, and how strongly these genetic instructions are utilized. Together these regions encode the information to direct the function and features of all living systems.

Genomes, as the product of billions of years of continuous evolution including the rewriting, rearrangement, addition, and removal of genomic elements^1^, are complex and dynamic networks. The intrinsic complexity of genomes presents a substantial barrier to completely understanding the full functionality of and the relationships between all genomic components. Even in *Escherichia coli* (*E. coli*)—the most extensively characterized model organism—the functions of 16% of genes remain fully uncharacterized^2^, and their regulatory mechanisms even less so^3^. The inability to fully elucidate the regulation, function and relationships of all genomic components in *E. coli* is driven by the intrinsic complexity and dynamic nature of its genome. These limitations can be even more pronounced for other, less studied and annotated, organisms.

Genome minimization provides an experimental approach for investigating this complexity by identifying and removing non-essential genomic segments, leaving behind only the essential genomic elements which are required to maintain a living system in a specific environment. In addition to deepening the understanding of natural genomes, a practical goal of many minimization efforts is to construct a genomic chassis of reduced size to serve as a modular platform upon which synthetic components can be added with reduced interference and complexity^4^. Reduced genomes also provide a more manageable platform for full genome synthesis and recoding efforts^5–9^.

Early, foundational genome minimization efforts relied heavily on prior information to inform reduced genome construction. In addition to regulation and function of individual genomic components, these studies probed gene essentiality using methods like single gene knockouts^10^, CRISPR interference^6^, and transposon mutagenesis^11^ to determine which genomic segments would be included in minimal genome designs. Although these methods excel in determining the essentiality of individual genes at the wild-type genome level, they struggle to address the unpredictability associated with combinatorial deletions^12^.

Historically, genome minimization efforts relied on predetermined information to inform the processes of either *de novo* reduced genome construction^5,13,14^ or genome reduction by sequential deletions^15–18^. Although the utilization of prior information for *de novo* reduced genome construction was used to produce the first synthetic reduced genome (*Mycoplasma mycoides* JCVI-Syn3.0)^5^, initial designs failed to result in a viable genome, resulting in the initiation of several design-build-test cycles before finally producing a viable genome. A similar effort in *E. coli* failed to yield viable cells^14^. Such workflows are often hindered by laborious construction and the risk of costly failure due to incomplete biological information. With this approach, a single design flaw or construction error could render the organism non-viable. (**Figure 1a**).

**Figure 1.**
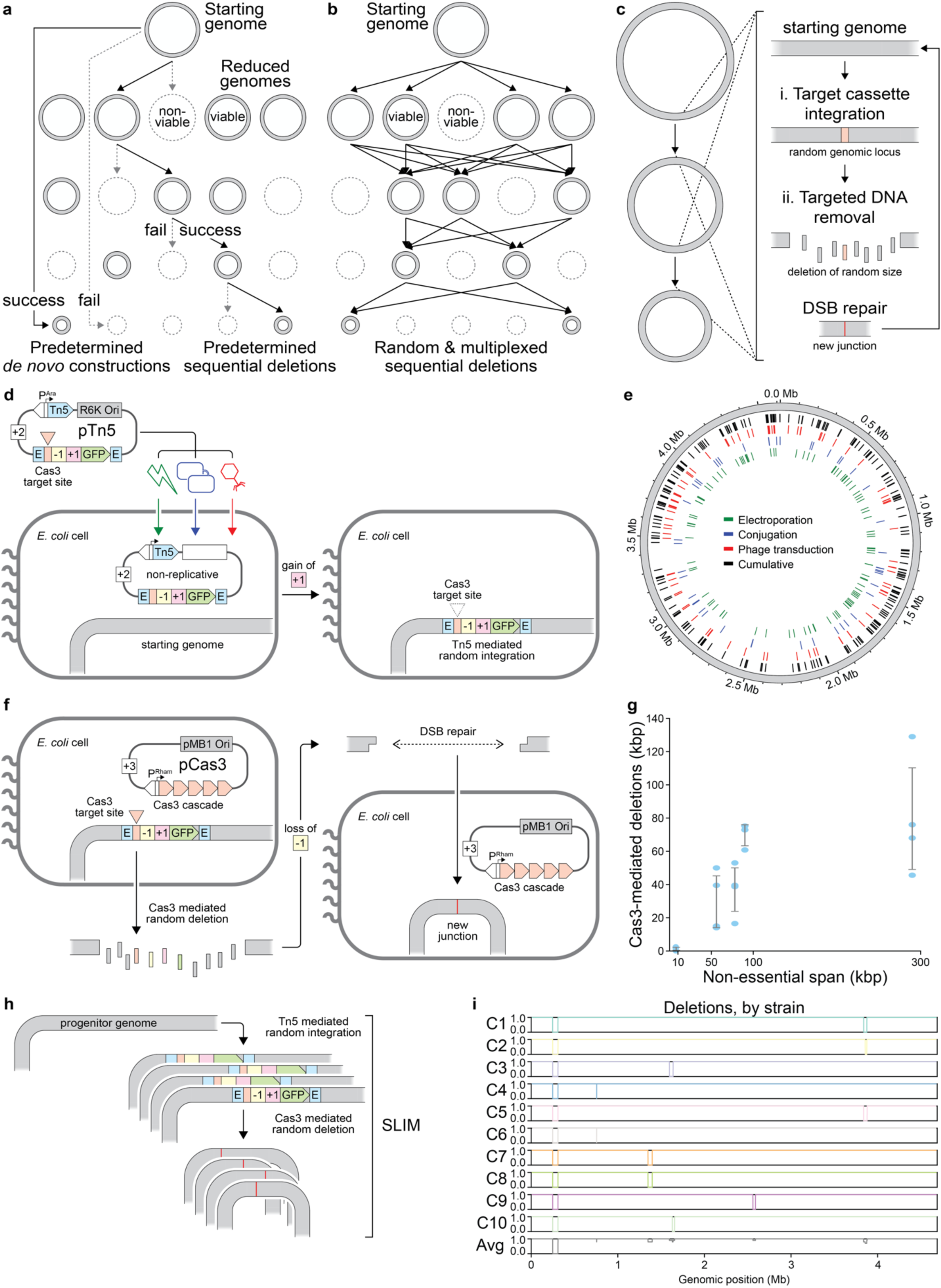
Conceptual diagrams and deconstructed workflow of SLIM. **(a)** Conceptual schematic demonstrating the workflow of traditional genome reduction methods. These methods rely on *a priori* information to make predetermined, sequential deletions or construct *de novo* reduced genomes. This approach can lead to the development of non-viable genomes, halting progress and resulting in costly delays and restarts. When successful, this approach often leads to just a single reduced genome. **(b)** Conceptual schematic demonstrating the workflow of SLIM-mediated genome reduction. This method hedges against the generation of non-viable genomes by making random, sequential deletions. When conducted on independent lineages, in parallel, this method will produce a library of genome-reduced lineages. **(c)** Schematic depicting the main steps of SLIM, which are repeated iteratively to generate a reduced genome: First, a target cassette is integrated into a random genomic locus, providing a nuclease target site. Then, a deletion of random size is made at and flanking the target site, resulting in the loss of the target cassette and surrounding DNA. Finally, the resulting DSB is repaired prior to the beginning of the next round. **(d)** Schematic depicting the random integration of the target cassette. Briefly, pTn5—a non-replicative vector—encoding a hyperactive Tn5 transposase and transposon cargo consisting of a Cas3 target cassette is delivered to pre-induced target cells using one of three methods: electroporation, conjugation or phage transduction. Once inside of target cells, Tn5 activates and transposes the target cassette from pTn5 to a random genomic locus. **(e)** Diversity of target cassette insertion sites following random target cassette integration, stratified by delivery method. Cumulative insertions are depicted in black. **(f)** Schematic depicting the DNA deletions mediated by Cas3 using SLIM. Following the random integration of the target cassette, expression of the Cas3-cascade is induced, prompting degradation of the target cassette and flanking genomic DNA. The resulting DSB is repaired prior to the onset of the next round of SLIM. **(g)** Evaluation of the size of deletions generated following Cas3-mediated DNA removal of target cassettes intentionally placed in 5 genomic loci, indicated by the distance between flanking essential genes (non-essential span). **(h)** Schematic representation of SLIM-mediated genome reduction in *E. coli*. The entire SLIM workflow was applied for a single round to the progenitor strain. **(i)** Coverage plot depicting the deletions generated following the application of SLIM to the progenitor strain. Ten colonies were selected for evaluation and deletions were identified following WGS.

An alternative approach, genome reduction by sequential deletions, proceeds by iteratively deleting pre-identified non-essential regions. Notable efforts generated strains such as MDS42 and DGF-298, the former achieving a 14.3% reduction through 42 rounds of deletions and the latter achieving a 37.4% reduction through 96 cumulative rounds of deletions^15–18^. However, the dependance of this approach on prior annotation of gene function, regulation, and essentiality makes it susceptible to previously unknown non-viable designs (**Figure 1a**) and limits its portability to other, less-studied organisms.

Building upon these strategies we propose an alternative approach, utilizing random and multiplexed sequential deletions (**Figure 1b**). Rather than relying on a single defined path (**Figure 1a**), here we rely on multiple random paths of sequential deletions attempted in parallel. The non-viable intermediates auto-eliminate themselves and the viable intermediates survive and propagate to the next round. Through iterative repetitions of this cycle, viable paths of genome reduction eventually emerge through the survival of viable intermediates (**Figure 1b**). This approach fundamentally eliminates the need to rely on any prior information regarding essentiality. Instead, it determines gene essentiality intrinsically.

Prior attempts at this approach include methods like RANDEL and TMRD^19,20^. While successful in principle, each of these methods takes at least 7 days to delete a single segment and cumulatively achieved only 2.6% to 5.5% reduction over 5 rounds of genome reduction^19,20^ The limited scope of these studies generated only a small number of genome-reduced *E. coli* strains with substantially overlapping deletions. In addition, the reliance of these methods on endogenous *E. coli* factors to remove randomly targeted genomic segments limits their cross-species portability.

To overcome these limitations, we developed Stochastic Lineage-based Iterative Minimization (SLIM). SLIM is a modular, high-throughput platform designed for unbiased DNA removal and compatibility across large phylogenetic distances. The foundational features of SLIM—the generation of multiple reduced genomes in parallel combined with the random nature of deletions—allow for survival-based selection of non-lethal deletions, eliminating the risk of lethal deletions that frequently affect traditional methods (**Figure 1b**).

SLIM transforms genome minimization from a single-strain pursuit to a systems-level exploration of the global bacterial genomic design space. In this study, we deploy SLIM in *E. coli* DH10B, a derivate of the *E. coli* K-12 lineage, for 21 iterative rounds of genome reduction, completing a single round in as little as 3 days and generating a library of 10 genome-reduced lineages (**Figure 2a-c**). Detailed phenotypic and molecular characterization of this library revealed new insights into deletion tolerance, the effects of deletions on expression dynamics, and context dependent phenotypic behaviors driven by expression dynamics (**Figures 2-4**). We further demonstrate the generality of SLIM by successfully deploying it in *Shigella flexneri* and *Pseudomonas putida* (**Figure 5**). Together, this work establishes SLIM as a scalable, unbiased, and generalizable platform for unlocking the landscape of reduced genomes, moving us closer to identifying the fundamental components of a minimized genomic chassis.

**Figure 2.**
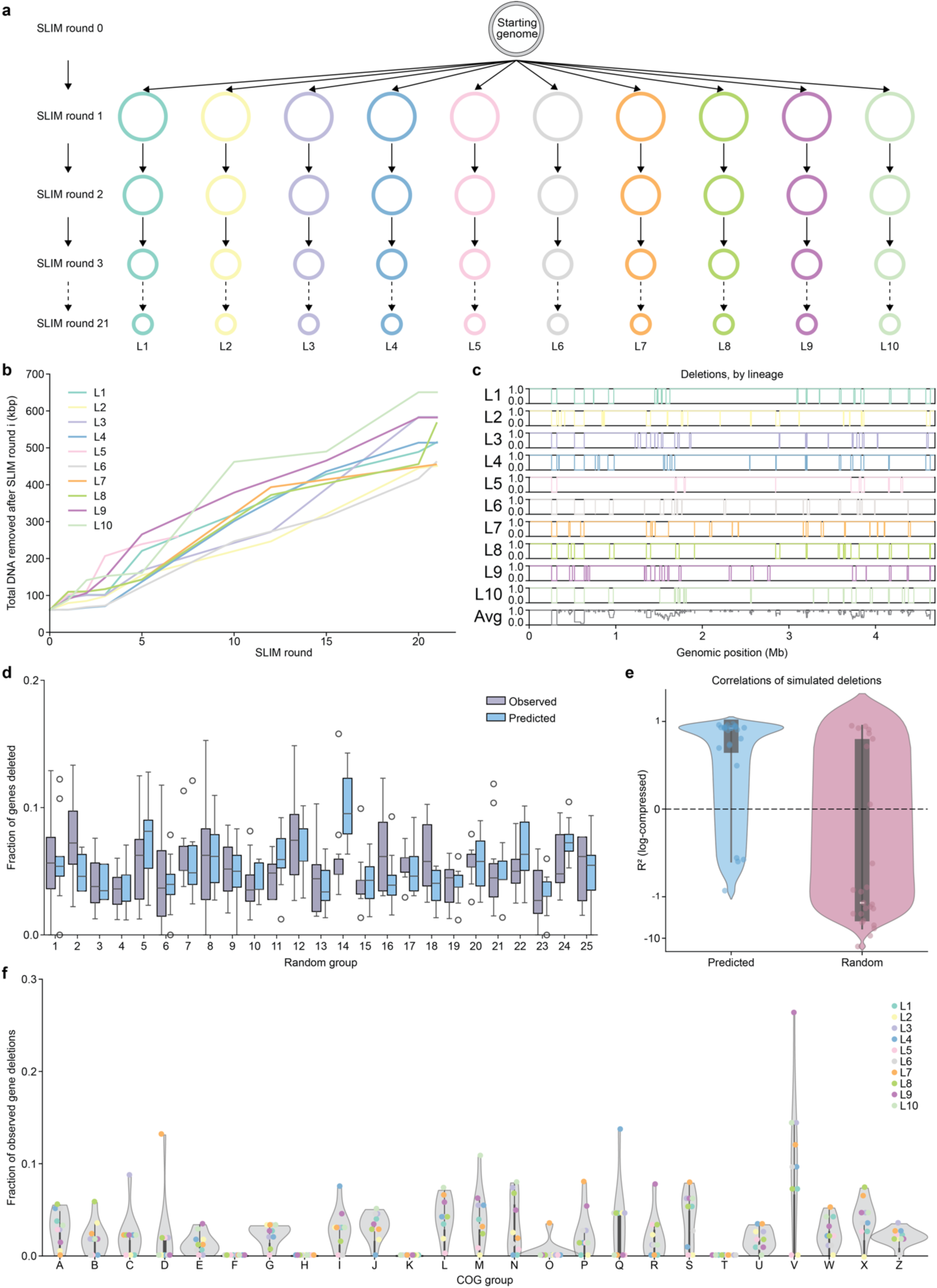
Evaluations of deletions generated following 21 rounds of SLIM-mediated genome reduction in *E. coli*. **(a)** Schematic representation of the 21 rounds of SLIM. Round 1 of SLIM was applied to a singular starting strain, generating 10 independent and distinct lineages. SLIM was applied to each lineage independently in successive rounds. **(b)** Total amount of DNA removed, by lineage, across 21 rounds of SLIM. **(c)** Coverage plot displaying the genomic location and size of every deletion identified, per lineage, across 21 rounds of SLIM. The “Average” row depicts the average coverage, per nucleotide, across all 10 lineages. **(d)** Fraction of observed deletions (lavender) and model-generated deletion predictions (light blue) for each of 25 random randomly generated groups following 21 rounds of SLIM. **(e)** Coefficient of determination values (R2) for deletion predictions generated by the genomic location-based deletion probability model (light blue) and the randomized deletion probability model (pink). **(f)** Violin plots depicting the fraction of genes deleted across 25 COG groups for each of the 10 genome-reduced strains. Plots were bounded on the y-axis by data 0 and the maximum data point. COG groups Key: [A] RNA processing and modification; [B] Chromatin structure and dynamics; [C] Energy production and conversion; [D] Cell cycle control, cell division, chromosome partitioning; [E] Amino acid transport and metabolism; [F] Nucleotide acid transport and metabolism; [G] Carbohydrate transport and metabolism; [H] Coenzyme transport and metabolism; [I] Lipid transport and metabolism; [J] Translation, ribosomal structure and biogenesis; [K] Transcription; [L] Replication, recombination and repair; [M] Cell wall/membrane/envelope biogenesis; [N] Cell motility; [O] Posttranslational modification, protein turnover, chaperones; [P] Inorganic ion transport and metabolism; [Q] Secondary metabolites biosynthesis, transport and catabolism; [R] General function prediction only; [S] Function unknown; [T] Signal transduction mechanisms; [U] Intracellular trafficking, secretion, and vesicular transport; [V] Defense mechanisms; [W] Extracellular structures; [X] Mobilome: prophages, transposons; [Z] Cytoskeleton.

**Figure 3.**
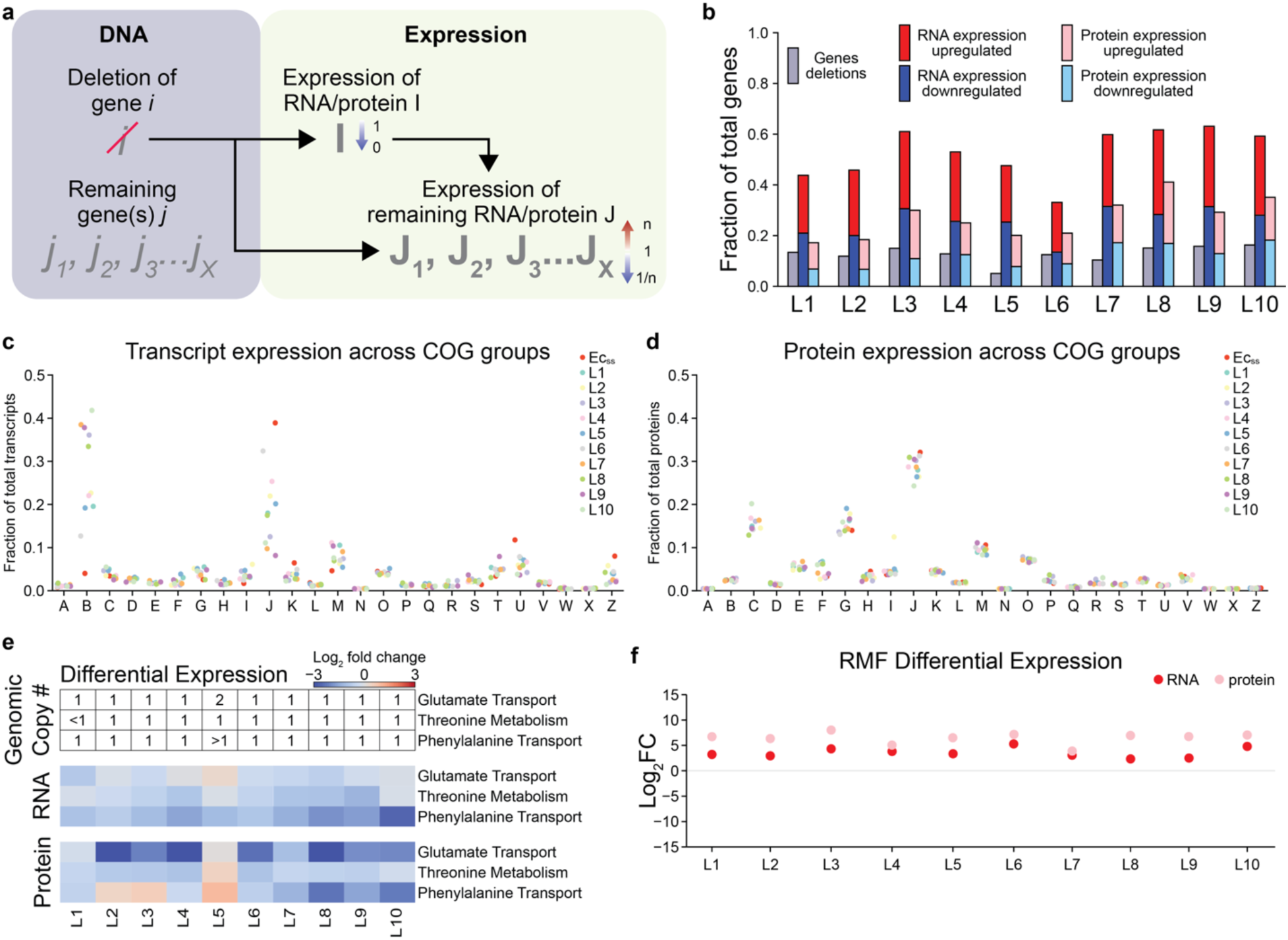
Evaluation of the expression dynamics of the remaining genome, following 21 rounds of SLIM-mediated genome reduction. **(a)** Schematic describing the anticipated effects on the expression of RNA and protein *I* and RNA and protein *J* (for *J_1_,…,J_X_*) following the deletion of gene *i*. **(b)** Bar plots depicting the fraction of total genes deleted (lavender), genes whose transcripts exhibit up- or down-regulation (red, blue) and genes whose protein products exhibit up- or down-regulation (pink, light blue). **(c)** Fraction of total transcripts, per lineage, attributed to each of 25 COG groups. **(d)** Fraction of total proteins, per lineage, attributed to each of 25 COG groups. **(e)** Differential transcript (top) and protein (bottom) expression for three pathways across all 10 lineages. The average genomic copy number following 21 rounds of SLIM for genes within each pathway is indicated, per genome-reduced strain. **(f)** Differential expression of RMF transcripts (red) and proteins (pink) across all 10 lineages.

**Figure 4.**
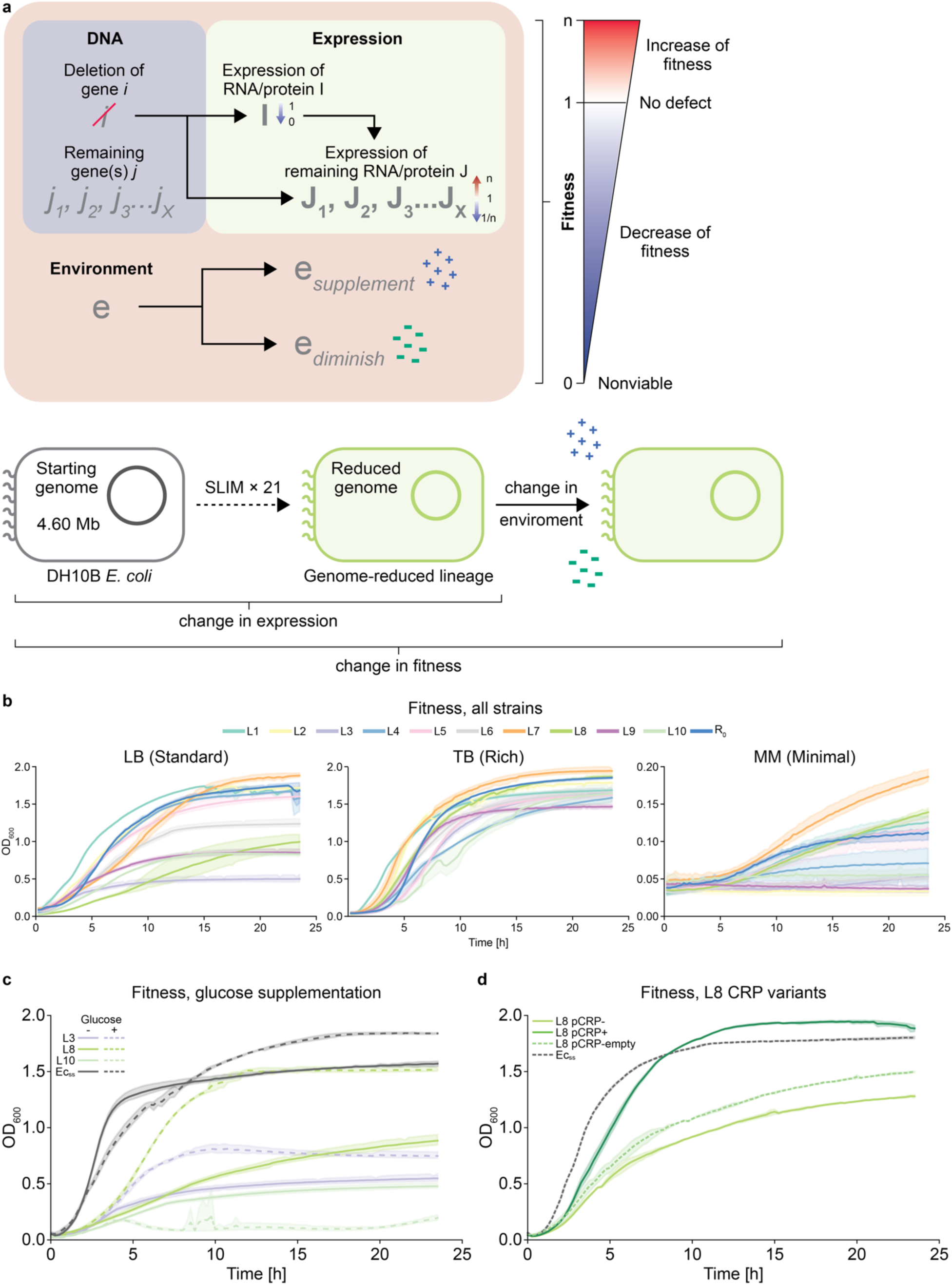
Evaluation of fitness changes due to expression changes and environmental perturbations. **(a)** Extension of Figure 3a, depicting the predicted effects on cellular fitness of environmental perturbations, in the context of nutrient supplementation (blue) or depletion (green), when combined with previously observed changes in expression. **(b)** Fitness curves for derivatives of genome-reduced *E. coli* lineages and the progenitor strain (R_0_) in three media spanning a gradient of nutrient capacity: standard (left), rich (middle), and minimal (right). **(c)** Fitness curves for three genome reduced strains—L3, L8, and L10—and Ec_ss_ when grown in standard media (LB) and standard media supplemented with glucose (LBg). **(d)** Fitness curves for Ec_ss_, L8, L8 + pCRP, and L8 + pCRP-empty (control vector encoding only the origin of replication and antibiotic resistance gene) when grown in LB.

**Figure 5.**
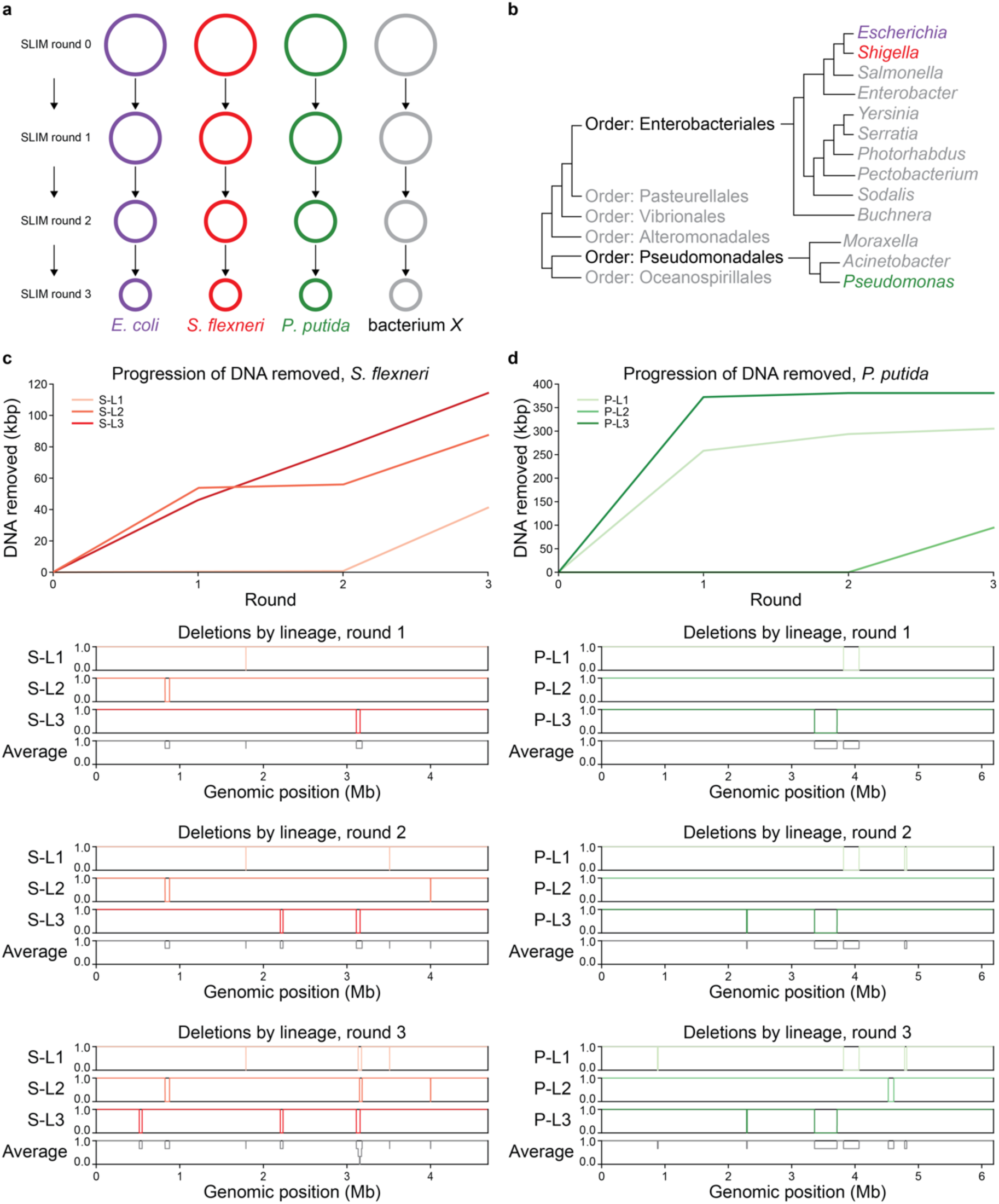
Utilization of SLIM to reduce the genomes of *S. flexneri* and *P. putida*. **(a)** Schematic representation of the application of SLIM to reduce the genomes of diverse bacteria. SLIM was used to reduce the genomes of *E. coli* (lavender), *S. flexneri* (red), and *P. putida* (green) and can be used to reduce the genomes of many other bacteria (gray). **(b)** Map exhibiting the phylogenetic distance between *E. coli* and the other bacteria to which SLIM was applied, *S. flexneri* and *P. putida*. **(c, top)** Total amount of DNA removed, by lineage, across 3 rounds of SLIM in *S. flexneri*. **(c, remaining)** Coverage plot displaying the genomic location and size of every deletion identified, per lineage, for each of 3 rounds of SLIM in *S. flexneri*. The “Average” row depicts the average coverage, per nucleotide, across all 3 lineages. **(d, top)** Total amount of DNA removed, by lineage, across 3 rounds of SLIM in *P. putida*. **(d, remaining)** Coverage plot displaying the genomic location and size of every deletion identified, per lineage, for each of 3 rounds of SLIM in *P. putida*. The “Average” row depicts the average coverage, per nucleotide, across all 3 lineages.

## Results

### Design and development of SLIM for scalable and unbiased genome reduction

SLIM operates iteratively to remove a contiguous genomic segment, of random size, from a random genomic locus through two sequential steps (**Figure 1c**). During the first step, a target cassette is inserted into the genome at a random genomic locus. During the second step, the target cassette and flanking endogenous DNA are degraded by a guided and progressive nuclease system. The amount of genomic DNA removed is random and unspecified; it varies from cell-to-cell and by genomic location. Due to the unspecified nature of the DNA removal process essential genes may be deleted in any individual cell, causing cell death. As a result, only tolerable deletions which result in the removal of only non-essential DNA propagate through successive rounds of genome reduction via viable intermediates. These two steps are repeated iteratively, each time shrinking the genome until all non-essential DNA has been removed, yielding a terminally reduced genome.

The random integration of the target cassette is initiated by the delivery of the pTn5 plasmid generated for this study. pTn5 encodes a hyperactive Tn5 transposase^21^ and a transposable target cassette. The target cassette is composed of a programmable target sequence that is recognized by the progressive nuclease system, a hygromycin resistance gene used as a positive selection marker (hygR, +1), a mutant allele of pheS^22^ used as a negative selection marker (pheS T251A_A294G, hereafter pheS^mut^, -1) for counterselection, and the GreenLantern variant of green fluorescent protein (GFP)^23^ used as an independent visual indicator (**Figure 1d**). Upon delivery of pTn5 to target cells, via electroporation, conjugation, or phage transduction, Tn5 transposase is activated immediately, resulting in the integration of the target cassette into a random genomic locus (**Figure 1d**). The pTn5 plasmid relies on the π-replicator (piR)/R6Kɣ system for replication^24^, but strains targeted for minimization using SLIM lack the piR gene. As a result, strains targeted for minimization initially lack the ability to replicate pTn5 and maintain associated components and features such as antibiotic (hygromycin) resistance. This design feature compels genomic integration of the target cassette ensuring the maintenance of hygR, leading to the enrichment of only cells which have randomly integrated the target cassette, upon antibiotic challenge.

We demonstrated and compared the efficacy and diversity of target cassette delivery and integration across the three distinct DNA delivery methods: electroporation, conjugation, and phage transduction. Each delivery method, when paired with Tn5 transposition, successfully facilitated broad distribution of cassette insertions across the genome (**Figure 1e**), yielding 86, 47, and 96 unique insertion sites out of approximately 200 isolated colonies, respective to each delivery method (**Figure 1e**).

Following the random integration of the target cassette, the target cassette and flanking genomic DNA are removed by the action of the CRISPR-Cas3 system encoded on the pCas3 plasmid. This system uses an RNA-guided multi-protein cascade complex and the Cas3 helicase-nuclease to identify and degrade DNA containing the programmed target sequence^25^. Upon induction of the Cas3-cascade, the cascade complex locates the target sequence within the target cassette then recruits Cas3 which progressively degrades the target cassette and flanking genomic DNA of random size (**Figure 1f**). Resulting double-stranded breaks are repaired by the cell’s endogenous DNA repair machinery. Unlike pTn5, pCas3 persists across multiple rounds of SLIM-mediated genome reduction. The components of pCas3 are conditionally activated upon induction.

In isolated experiments, Cas3-mediated DNA removal resulted in deletions consistent with the combined distance from the random integration site to the nearest flanking essential genes, a natural barrier to deletions, defined as “non-essential span”. DNA removal was found to be dependent on the negative selection pressure mediated by the negative marker pheS^mut^ (**Methods**). Initial DNA removal experiments utilized an earlier version of the target cassette (original target cassette); composed of a target sequence, hygR, and GFP, but without pheS^mut^. In the absence of negative selection, driven by pheS^mut^, Cas3-mediated DNA removal attempted on five differently sized (8 kbp, 56 kbp, 78 kbp, 90 kbp, and 290 kbp) non-essential spans yielded few to no GFP^-^ colonies (**Extended Data Fig. 1a**). In experiments using an updated design of the target cassette, with negative selection enabled by pheS^mut^ incorporation the target cassette (modified target cassette), successful Cas3-mediated target cassette degradation was achieved across all five non-essential spans (**Extended Data Fig. 1a**), with two of the five yielding degradation efficiencies above 90%. Whole genome sequencing of colonies which exhibited Cas3-mediated degradation revealed that deletions sustained from within the same target site were of different sizes, but deletion sizes varied with the sizes of non-essential spans, ranging from a 0.35 kbp deletion in the 8 kbp span to a 129 kbp deletion in the 290 kbp span (**Figure 1g**). A strain generated from targeting the 90 kbp span, which exhibited approximately 60 kbp of DNA loss, was chosen as the progenitor strain (R_0_), to which SLIM was applied for subsequent genome reduction (**Extended Data Fig. 1b**).

By combining Tn5-mediated random target cassette insertions with randomly sized deletions mediated by Cas3, we composed the full SLIM cycle. We then applied SLIM to R_0_, yielding ten genome-reduced strains (**Figure 1h**). The genomic locations of these deletions collectively span over 3 Mbp of the *E. coli* genome, from the 0.756 Mb locus to the 3.879 Mb locus (**Figure 1i**). Deletions in colonies 1, 2, and 5 exemplify the random nature of Cas3-mediated DNA removal. These deletions overlap within the same general genomic region yet differ in size substantially, 36 kbp, 18.6 kbp, and 40.8 kbp, respectively (**Figure 1i**). This result demonstrates that distinct Cas3 degradation events can converge on overlapping regions while producing unique deletion outcomes, further underscoring the stochastic and flexible nature of the SLIM system.

### The utilization of SLIM to generate a library of genome-reduced *E. coli* lineages

Four variations of the SLIM workflow, each developed to decrease the amount of time required per round of SLIM, were utilized across 21 rounds of genome reduction, ultimately generating a library of 10 unique genome-reduced *E. coli* lineages from R_0_, hereby referred to as L1 – L10 (**Figure 2a**). While Tn5 transposition and Cas3-mediated DNA removal parameters remained largely consistent, we optimized pTn5 delivery methods and timings across rounds (**Extended Data Fig. 1c-e**). For rounds 1–3 electroporation was used for delivery of pTn5. When utilizing electroporation as the pTn5 delivery method, each round of SLIM took 5 days to complete. For rounds 4–16 conjugation was used for pTn5 delivery, a change which dropped the time per round from 5 days to 4 days. Phage transduction was used for pTn5 delivery for rounds 17-21. For round 21, additional workflow optimizations dropped the time per round from 4 days to 3 days (**Methods**).

Across 21 rounds of SLIM, we observed 16 to 19 deletions per lineage, with deletion sizes ranging from 75 bp to 170 kbp (**Extended Data Table 1**). The cumulative DNA removed per strain ranged from 434 kbp to 649 kbp, representing a 9.25% to 13.9% reduction of the starting strain’s, *E. coli* DH10B (Ec_ss_), genome (**Figure 2b**). Deleted segments were dispersed throughout the entire genome. Aside from the original deletion in R_0_, there were no shared deletion events across all ten lineages (**Figure 2c**). Derivatives of nine out of the ten lineages had lost at least part of one of the two 113 kbp tandem repeat regions found in the Ec_ss_ genome, a previously described genomic feature^26^. Additional regions with high or low deletion frequency were observed as evidenced in **Figure 2c** (‘Average’ row).

### Investigating the tolerability of SLIM-mediated deletions

The intrinsic stochasticity of SLIM-mediated deletions ensures that only tolerable deletions are captured, allowing us to bypass impediments such as quasi-essential genes and combinatorial lethal gene pairs^4,5,27,28^. As a result, we are free to explore the full range of possible deletions to probe the deletion tolerance of individual genes. Comprehensive analyses revealed that the deletion tolerance of any individual gene is driven by a combination of factors, including genomic location, function, and genomic neighborhood. The contributions of these factors to deletion tolerance were examined through whole genome sequencing analyses*, in silico* deletion simulations, and transcriptomic and proteomic expression analyses.

To understand the role that genomic location plays in the determination of deletion tolerance, we conducted 10 *in silico* pan-genome deletion simulations (**Methods**) using only the genomic location of previously observed deletions and the frequency of these deletions across genome-reduced strains as prior information. After sorting genes into 25 random groups, we quantified the proportion of deleted genes per group and compared these deletion predictions to observed deletions (**Figure 2d**).

The model generated using computed deletion probabilities substantially outperformed a parallel model generated using random deletion probabilities, according to the coefficient of determination (R^2^, **Figure 2e, light blue**). R² values varied but were generally positive, indicating the meaningful predictive power of genomic location for determination of deletion tolerance. For comparison, we randomized the deletion probabilities across bins while maintaining group assignments and performed 10 additional simulations. These randomized controls yielded substantially lower or negative R² values (**Figure 2e, red**). Overall, average R² values were 0.47 for computed deletion probability-based predictions vs. –2.2 for randomized deletion probability-based predictions, indicating that our model utilizing genomic location-based deletion probabilities captures roughly half of the observed variance in deletions data. This suggests that the probability that a deletion can be tolerated correlates moderately with its specific position on the genome.

The role of gene function, described according to the 25 Clusters of Orthologous Genes (COG) functional categories^29^, in the determination of deletion tolerance varies by COG group (**Figure 2f**). Qualitative inspection revealed that certain categories, such as ‘Inorganic ion transport and metabolism (P),’ ‘Transcription (K),’ and ‘Secondary metabolite biosynthesis (Q),’ showed high levels of deletion tolerance. In contrast, categories such as ‘Translation, ribosomal structure, and biogenesis (J)’ and ‘DNA replication, recombination, and repair (L)’ exhibited markedly lower deletion frequencies (**Figure 2f**). This trend was further supported by analyzing the median deletion percentages per COG group (**Extended Data Fig. 2a**). While some groups either appear recalcitrant to deletions or were highly permissive, most fell within a moderate range, suggesting that gene function plays a non-deterministic role in deletion tolerance.

Features of genomic neighborhoods, such as toxin/antitoxin (TA) pairs and essential genes, demarcate possible deletion boundaries, as the deletion of such genomic elements often results in cell death. Using three independent essential gene datasets^10,11,30^ we compiled a list of consensus essential genes, deemed essential in all three studies, and additional genes deemed essential in at least one of the studies^10,11,30^ (**Extended Data Table 2**). Across 21 rounds of SLIM and 10 genome-reduced lineages, no consensus essential genes were deleted. An average of only 3.9 non-consensus essential genes were deleted from derivatives of each lineage. Analyses of TA pairs revealed intriguing bounding behavior. Within a 30kbp segment of the genome (loci 1.664-1.694 Mb), a cluster of deletions from derivatives L3,4,5,6,8 and 10 occupy over 28.3 kbp. The remaining 1.7 kbp is composed entirely of a segment encoding the hipA/B TA system (**Extended Data Fig. 2b**).

These findings illuminate the reality that deletion tolerance is highly complex and non-deterministic, influenced to varying extents by a combination of factors including, essentiality status, genomic location, gene function, and TA pairs. Together, these results demonstrate that a multi-lineage, library-based approach enables not only the generation of diverse genome-reduced strains but also a richer understanding of the forces and constraints that govern genome architecture.

### Deletions have a profound effect on the expression of remaining genomic components

As a result of the complex interdependencies amongst genes in the *E. coli* genome, deletion of a single gene can have extensive downstream effects on the remaining genomic components. While the deletion of an individual endogenous gene (‘*gene i*’) predictably eliminates the expression of RNA and protein *I*, the broader cellular consequences are often nontrivial (**Figure 3a**). The loss of both RNA and protein *I* can alter the expression levels of many genes (*j₁, j₂, …, jₙ*), triggering broad transcriptional and translational shifts (**Figure 3a**).

To investigate this, we performed both RNA sequencing and protein mass spectrometry on derivatives of all ten genome-reduced lineages after 21 rounds of SLIM, as well as on a non-reduced control strain (Ec_ss_, *E. coli* DH10B + pCas3). Multi-omics analyses reveal that across all derivatives of genome-reduced lineages, RNA expression changes were observed in 33.1%–63.1% of remaining genes and protein expression changes in 17.2%–41.1%, despite only 5.1%–16.3% of genes being deleted (**Figure 3b**). Both RNA and protein differential expression (DE) analyses were performed, comparing each individual genome-reduced derivative to Ec_ss_. Volcano plots summarizing DE results, including indication of the top five up- and downregulated transcripts/proteins for each strain, are provided in **Extended Data Figs. 3 and 4**.

Transcriptomic and proteomic analyses conducted across 25 COG groups revealed the extent of expression changes resulting from deletions sustained during the genome reduction process for derivatives of genome-reduced lineages as compared to Ec_ss_. Transcriptome-level expression assessments revealed that derivatives of genome-reduced lineages allocate a larger fraction of RNA expression to genes involved in ‘Chromatin structure and dynamics (B)’ compared to Ec_ss_ (**Figure 3c**), suggesting an emphasis on DNA protection via genes such as hns (mean log_2_FC = 3.276)^31^, dps (4.084)^32^, and hupA (1.015)^33^ This analysis also revealed that Ec_ss_ devotes over 40% of its transcriptomic output to ‘Translation, ribosomal structure, and biogenesis (J),’ a category consistently downregulated, in some cases substantially, across genome-reduced lineages. This trend persists at the proteomic level, albeit less dramatically (**Figure 3d**). Variations among derivatives of genome-reduced lineages were also seen in ‘Intracellular trafficking (U),’ ‘Cell wall/membrane/envelope biogenesis (M),’ and other categories. By contrast, all strains, including Ec_ss_, show minimal and stable expression in categories like ‘Defense mechanisms (V),’ ‘RNA processing and modification (A),’ and ‘Cell motility (N).’

Analyses of pathway-level DE data highlighted pathways of interest, revealing potential gaps in pathway annotations and a novel counterselection escape mechanism. To explore changes in specific pathways, we compared RNA and protein DE values for each strain versus Ec_ss_. Amino acid transport and metabolism pathways, additional metabolic pathways, and stress response pathways were evaluated (**Figure 3e, Extended Data Fig. 5a**). Glutamate transport and threonine metabolism pathways exhibited a pattern of downregulation in all derivatives except L5 across RNA and protein DE analyses (**Figure 3e**). In the case of glutamate transport—largely mediated by multiple genes (gltI, gltJ, gltK, gltL, gltS)^34–36^—this trend is likely explained by the genomic location of these genes in the tandem repeat region of the Ec_ss_ genome. As L5 retains both copies of the tandem repeat segment, the genomic copy number for gltI, gltJ, gltK, gltL, and gltS remains at 2. In all other genome-reduced strains, one copy of the tandem repeat segment has been deleted, reducing the genomic copy number of gltI, gltJ, gltK, gltL, and gltS to 1 in these strains (**Figure 3e**). Genes regulating threonine metabolism, (including thrA, thrB, thrC, thrL, and tdcA, tdcB, tdcC, tdcD, tdcE, tdcF, tdcG) ^34–36^, are not located in this repeat region. Furthermore, the genomic copy number of these genes remains at 1 across all genome-reduced derivatives except for L1 (tdcD, tdcE, tdcF, and tdcG deleted). Yet we observe a systematic downregulation of this pathway which mirrors that of glutamate transport (**Figure 3e**). This pattern of expression, in concert with the deletion of one half of the tandem repeat segment that is shared by L1-4 and L6-10 which results in a decrease in copy number for genes in this region, suggests the possible presence of an uncharacterized regulator within the deleted tandem repeat segment.

While the copy number of genes from the phenylalanine transport pathway, required for import of 4-chlorophenylalanine and full functionality of the pheS^mut^ counterselection strategy, is largely unchanged across all genome-reduced derivatives, transcriptomic expression levels are globally downregulated (**Figure 3e**). Interestingly, protein expression levels for this pathway fluctuate across all genome-reduced derivatives (**Figure 3e**). As SLIM was designed specifically to prevent SNP-mediated escape from pheS^mut^ counterselection through the reintroduction of the pheS^mut^ gene at the start of each round of SLIM, the downregulation of the phenylalanine transport pathway represents a novel and unexpected counterselection escape mechanism. These data help to illuminate the complex global genomic reprogramming that takes place during the genome reduction process.

Amongst other pathways of interest, hydrogen peroxide (H₂O₂) stress response and starvation stress response exhibited consistent upregulation across derivatives of genome-reduced lineages (**Extended Data Fig. 5a**), suggesting a trend towards adaptation to oxidative stress and metabolic deficiency conditions.

Some pathways, such as cysteine transport in derivatives L6–8 and auto-inducer II (AI2)-mediated quorum sensing in L5 (just RNA) and L8 (both RNA and protein), sustained substantial deletions, yet genes which remained intact exhibited consisted upregulation (**Extended Data Fig. 5a**). These findings highlight a key distinction: while whole pathways may be affected by deletion events, surviving genes may undergo compensatory regulation. Conversely, some pathways exhibit consistent regulation across all genome-reduced strains: glutamine transport and histidine metabolism are generally upregulated, while alanine and phenylalanine transport and phosphonate/phosphinate metabolism are consistently downregulated.

Finally, one especially striking observation emerged from single gene DE analyses: all genome-reduced strains exhibited substantial upregulation of RMF, encoding the ribosome modulation factor (**Figures 3f**). RMF promotes ribosome hibernation and stabilization by sequestering ribosomal subunits as the cell prepares to enter stationary phase^37^. This is consistent with a general shift in expression strategy and resource conservation amongst derivatives of genome-reduced lineages.

### Examining the interplay between environmental context and expression changes

The effects of the fitness-altering expression changes induced by deletions can be exacerbated or alleviated by changes to the environment via nutrient supplementation or deprivation (**Figure 4a**). After 21 rounds of SLIM-mediated genome reduction, we assessed the fitness of derivatives of all ten genome-reduced lineages compared to Ec_ss_ (**Extended Data Fig. 5b**) and R_0_ (**Figure 4b**) in standard media (Luria-Bertani medium, LB), rich media (Terrific broth, TB) and minimal media (M9 mineral medium, MM)^38^. Six derivatives (L1, L2, and L4–L7) exhibited growth behaviors similar to R_0_, with minor differences in lag phase, doubling time, and saturation density in standard media (**Figure 4b**). In contrast, L3, L8, L9 and L10 exhibited longer doubling times and substantially reduced saturation densities in standard media. To explore the compounded effect of changes in expression with substantial nutrient supplementation we assessed the fitness of derivatives of genome-reduced lineages in rich media. In rich media, all derivatives exhibited a fitness level comparable to R_0_ (**Figure 4b**). The growth deficits observed for L3, L8, L9 and L10 in standard media were largely alleviated. Next, we explored the effect of changes in expression combined with substantial nutrient deprivation through fitness assessments in minimal media. In minimal media several derivatives exhibited no growth—L2, L3, L6, L9 and L10—while L1, L5 and L8 grew similarly to R_0_ (**Figure 4b**). Amongst all amino acids, minimal media supplies only leucine; a requirement for the growth of any strains derived from the genome reduction parental strain, Ec_ss_, due to a previously described deletion^26^. A single genome-reduced derivative, L7 (**Δ**455kb, ∼10%), exhibits improved fitness as compared R_0_, growing to a higher saturation density in all three media. L7 exhibits comparable fitness to Ec_ss_ in standard media and rich media but is substantially less fit in minimal media (**Extended Data Fig. 5b**).

In addition to overall compositions of the media, a simple media supplementation also elicited changes in fitness for a subset of derivatives of genome-reduced lineages. The addition of a single nutrient, glucose, to standard media (LBg), affected the fitness of three derivatives, L3, L8 and L10, in different and sometimes substantial ways (**Figure 4c**). Experiments conducted in standard media supplemented with glucose revealed a marginal increase in fitness for L3 and a surprising decrease in fitness for L10, as compared to growth in standard media. L8, however, exhibited a substantial increase in fitness, exhibiting a substantial decrease in doubling time during exponential phase and reaching a saturation density comparable to Ec_ss_, when grown in standard media supplemented with glucose as compared to standard media without glucose.

An analysis of 385 pathways containing genes regulated by glucose via the cyclic-AMP(cAMP)-CRP intermediate^39^ revealed plausible drivers behind the observed fitness change for L8 in standard media supplemented with glucose as compared to standard media. Transcriptomic and proteomic analyses revealed downregulation of many cAMP-CRP associated pathways in L8, including those responsible for processing critical amino acids such as tryptophan, threonine, alanine, serine, glutamate and valine (**Extended Data Figs. 6 and 7**). These analyses also revealed significant overexpression of RspAB in L8 and other genome-reduced strains (**Extended Data Fig. 8a-b**), regulators of stationary phase that are indirectly repressed by cAMP-CRP and promote growth arrest^40^. Principal component analysis (PCA) of whole-pathway transcriptomic and proteomic data for all cAMP-CRP associated pathways across all genome-reduced strains revealed a noticeable separation for L8 across two principal components (**Extended Data Fig. 8c-d**), indicating a coordinated shift in expression across several cAMP-CRP regulated pathways. Interestingly, PCA of transcriptomic data revealed separation across three principal components for both L3 and L10 as well (**Extended Data Fig. 8c**). Three-dimensional PCA of proteomic data revealed robust separation of L8 from the rest of the genome reduced strains (**Extended Data Fig. 8e-f**). These findings point to systematic dysregulation of cAMP-CRP regulated pathways for these strains, especially L8, consistent with their lack of fitness when grown in standard media. We propose the hypothesis, for L8 specifically, that supplementation of standard media with glucose likely alleviates this defect by bypassing CRP-mediated signaling.

Further examination of genomic deletions sustained by L8 revealed a deletion presumed to be the driver of the attenuated fitness of L8 in standard media. The deleted segment (loci 3.572 to 3.589 Mb) contains the CRP gene, the master regulator of the cAMP-CRP axis. To verify that this deletion was driving the observed fitness changes for L8 in standard media supplemented with glucose vs. standard media without glucose, we transformed L8 with a low-copy plasmid encoding the CRP gene while retaining endogenous CRP regulatory elements. Upon reintroduction of the CRP gene, we observe an almost complete recovery of fitness for L8 in standard media, as compared to Ec_ss_ (**Figure 4d**).

All together, these findings demonstrate the precarious interplay between environmental conditions and genome reduction. Deletion of endogenous DNA dramatically alters the expression patterns of remaining genes and changes in expression directly affect cellular fitness. However, expression-mediated fitness changes are highly context-dependent and can be reversed or exacerbated by environmental perturbations. These results emphasize the need to consider environmental context when engineering or utilizing genome-reduced organisms.

### SLIM: A modular method for genome reduction of diverse bacteria

The long-term ambition of the general genome minimization field is not limited to refining the genome of a single species; instead, the broader vision involves constructing truly novel organisms, which requires identifying the fundamental genomic elements of a wide array of phylogenetically distinct microbes. Achieving this goal demands the development and deployment of genome reduction methods that are generalizable across species (**Figure 5a**). To our knowledge no previously published genome reduction platform has demonstrated capabilities beyond a single organism.

The modularity and broad applicability of SLIM was demonstrated through deployment in two additional bacteria spanning different phylogenetic orders: *S. flexneri* and *P. putida* (**Figure 5b**). In *S. flexneri*, we used the standard SLIM workflow to generate a library of three genome-reduced lineages across three iterative rounds. After the third round, derivatives from each lineage were selected for whole genome sequencing. Each derivative sustained three deletions, ranging in size from 298 bp to 53 kbp (**Extended Data Table 1**), with total DNA removed per strain ranging from 37 kbp to 112 kbp (**Figure 5c**). For genome reduction of *P. putida*, we utilized a variation of the SLIM workflow that does not rely on pheS^mut^/4-chlorophenylalanine mediated counterselection. While this approach is not viable in *E. coli* (**Extended Data Fig. 1a**), the high activity of the Cas3-Cascade system in *Pseudomonas* species enabled efficient deletions without counterselection. After three rounds of SLIM in *P. putida*, an average of two deletions per strain were observed, ranging from 8.2 kbp to 356 kbp. The total amount of DNA removed from each derivative ranged from 88.4 kbp to 365 kbp (**Figure 5d**), with the latter representing nearly 6% of the total *P. putida* genome.

Importantly, because SLIM had already been fully developed and optimized in *E. coli*, deployment in *S. flexneri* and *P. putida* was straightforward, rapid, and robust, with genome reduction of each completed in parallel in less than one month. This demonstrates SLIM’s versatility and efficiency, revealing the potential of SLIM as a broadly applicable platform for unbiased genome reduction across diverse bacterial taxa.

## Discussion

SLIM transforms genome reduction from a slow, single-strain pursuit into a high-throughput, systems-level exploration. By generating diverse libraries of genome-reduced lineages rather than isolated strains, we provide the analytical power necessary to distinguish between universal biological responses and strain-specific adaptations. Landmark studies have historically required about a decade to produce a single minimized strain^5,15^. We showed that a single round of SLIM can be completed in as little as 3 days and were able to generate a library of genome-reduced *E. coli* in 21 rounds of SLIM, with genomes reduced by up to 14%. In *P. putida* we demonstrated the removal of almost 6% of the genome in 3 rounds of SLIM. This efficiency is a direct result of SLIM’s information-agnostic and multiplexed genome reduction approach; by remaining unbiased by prior essentiality annotations, which are often dynamic and context-dependent, SLIM avoids the non-viable dead ends that have negatively affected other genome reduction methods. By generating many reduced-genome strains in parallel, SLIM-mediated genome reduction can capture a wider breadth of deletions than traditional methods.

Our findings suggest that the tolerance of individual gene deletions is a multi-factorial property mediated by genomic location, gene function, and local genomic neighborhood. Analyses examining the contribution of gene function to deletion tolerance yielded mixed results. While our genomic coordinate-based model captured a portion of the observed deletion variance (R^2^ = 0.47), it remains insufficient to predict the full scope of deletions. The presence of essential genes and toxin-antitoxin pairs flanking deletions suggests that genomic geography may constrain the limit of genome reduction. Other factors that affect expression and may indirectly contribute to deletion tolerance include proximity to the origin of replication and colocalization of distant genomic regions due to the 3D structure of the genome^41,42^. These results indicate that, like the relationships between genes, the deletion tolerance of individual genes is complex and dynamic. As the genome reduction process progresses it is likely that the deletion tolerance of individual genes will adjust in unpredictable ways, underscoring the difficulty of *de novo* genome design and validating the need for empirical, stochastic methods like SLIM to map the boundaries of minimal genomes.

A primary insight from our multi-omics analysis is that derivatives of genome-reduced lineages became physiologically reprogrammed entities, substantially distinct from their wild-type ancestor. Previous studies have shown that deleting even a single gene can alter the expression of many others^43,44^. Our analyses indicate that the number of transcripts and proteins exhibiting differential expression far exceeded the number of deleted genes. We observed a persistent stress response evidenced by the global overexpression of RMF, typically expressed upon entrance into stationary phase^37^, and the downregulation of some standard metabolic functions. Divergences in expression patterns across derivatives of genome-reduced lineages, when combined with deletions data, demonstrate how genome reduction can expose gaps in current functional annotations.

Remarkably, DE analyses revealed a novel mechanism of counterselection escape. Rather than mutating the pheS^mut^ gene to relieve pressure from the pheS^mut^/4-chlorophenylalanine negative selection utilized during the DNA removal process, derivatives consistently downregulated the phenylalanine transport pathway. This systemic rerouting of expression to circumvent toxicity highlights the flexibility of bacterial adaptive landscapes and suggests that long-term selection pressure in minimized strains can drive unexpected cellular rewiring.

Environmental perturbation experiments revealed that the essentiality of a gene is often a function of the environmental niche. The restoration of fitness in the CRP gene deletion derivative L8, when grown in standard media supplemented with glucose, illustrates that the functional consequences of a deletion are not absolute, but are dictated by the interplay between the remaining genome and the environment.

The broadly spanning utility of SLIM is highlighted by the genome reduction of multiple, phylogenetically distant, bacteria. SLIM’s species-agnostic architecture allowed for the rapid minimization of *S. flexneri* and *P. putida*, remarkably, without organism-specific optimization. This modularity suggests that the SLIM platform can be readily adopted across the microbiology community to identify the fundamental building blocks of diverse bacterial taxa. As the field advances beyond bacteria, the principles established by SLIM—parallelization and unbiased DNA removal—can work synergistically with tools like SCRaMbLE^45^ in yeast or prime editing tools like PEDAR^46^ and PRIME-del^47^ in mammalian cells to accelerate the scope of genome reduction in more complex systems.

Altogether, this work highlights the analytical power and practical promise of utilizing SLIM to generate diverse libraries of genome-reduced lineages. The space of all possible minimized genomes is vast: for any given species, there exists a substantial number of viable reduced genome configurations, each reflecting a unique sequence of deletions and adaptive responses. Expanding into this space is essential for uncovering the fundamental building blocks of cellular life. Until now, the generation of genome-reduced strains has been a slow and manually intensive process, often taking years and yielding a limited view of this broader landscape. SLIM provides the first scalable platform to navigate this landscape, moving us closer to the rational design of modular genomic chassis and the eventual *de novo* synthesis of fully engineered organisms.

## Methods

### General methods

Unless otherwise stated, DNA was amplified using PrimeSTAR GXL (TaKaRa, #R050). Oligonucleotides were sourced from IDT and Millipore Sigma. Nucleic acid processing utilized QIAquick PCR Purification (QIAGEN, #28106), GeneJET Gel Extraction (Thermo Scientific, #K0691), and QIAprep Spin Miniprep (QIAGEN, #27106) kits. Plasmids were assembled via Gibson assembly (NEB, #E2621) and cloned into electrocompetent *E. coli* DH10B. Antibiotic/counter-selection concentrations: streptomycin (100 μg/ml), hygromycin (250 μg/ml), 4-chlorophenylalanine (5 mM), chloramphenicol (25 μg/ml), sucrose (7.5% w/v), kanamycin (50 μg/ml), carbenicillin (100 μg/ml), and gentamicin (15 μg/ml). All bacteria were cultured at 37°C with standard media (LB) and requisite antibiotics unless otherwise stated.

### Strains and plasmids construction

Briefly, parental strains included *Escherichia coli* DH10B rpsL K43R, *Pseudomonas putida* KT2440, and *Shigella flexneri* CFS100 rpsL K43R ΔrecA Δtus 1201 ΔoriT ΔhigB Δhok. *S. flexneri* was kindly provided by Shelley M Payne, University of Texas at Austin, as a gift. All other bacterial strains were acquired from ATCC. *E. coli* p1kc_piR-sC was constructed as described previously^48^. *E. coli* piR_genRK24, developed from the high frequency recombination *E. coli* DH10B_genRK24 strain (unpublished), was used as a donor strain to deliver pTn5-v5 via conjugation. Briefly, DH10B_genRK24 was generated by integrating elements from the pRK24 conjugative plasmid^52^ into the genome of *E. coli* DH10B. To engineer *E. coli* piR_genRK24, the piR-sC cassette was amplified and integrated into the *E. coli* DH10B_genRK24 genome. *E. coli DH10B*, *S. flexneri* and *P. putida* were all transformed with pCas prior to the application of SLIM for genome reduction.

Components used to construct pTn5-v2, v5 and v11 were taken from psfTn5 (Addgene #79107), Marionette cassette^49^, BBa_J72214-BBa_J72090 (Addgene #40782), and in-house plasmids. pCas3 was constructed from pCas3cRh (Addgene #133773); pdeL specific crRNA was amplified from the *E. coli* DH10B genome and cloned into pCas3 following digestion with BsaI (NEB, #R3733L). An inducible T4 ligase from pCA24N-ligase (Addgene #87741) was added to pCas3 in a sequential assembly step. For the pCRP plasmid, crp with native regulatory elements was amplified from the genome of *E. coli* DH10B and assembled into the low-copy pCRP-empty control vector, containing the pBelo BAC origin of replication and an Ampicillin resistance gene.

### Cas3-mediated DNA removal by non-essential span

Ten independent strains were generated to test the DNA removal efficacy of the Cas3-cascade system by integrating either the original target cassette (target sequence, hygR and GFP) or the modified target cassette (target sequence, pheS^mut^, hygR and GFP) into one of five genomic locations via lambda RED recombination^50^ encoded by the previously constructed recombination plasmid, pKW20. The five loci, indicated by non-essential span, are as follows: 8 kbp, 56 kbp, 78 kbp, 90kbp and 290 kbp. pKW20 was cured after integration verification then pCas3 was transformed.

Strains containing pCas3 and target cassettes were grown overnight in 3 mL LB + hygromycin and gentamicin. Overnight cultures were harvested (4000g, 6 min) and resuspended in LB containing gentamicin and 0.3% rhamnose for Cas3 induction. Cultures were incubated for 5.5 hours, then plated on LB agar plates supplemented with just gentamicin (original target cassette) or gentamicin + 4-chlorophenylalanine (modified target cassette) and incubated overnight. The ratio of GFP^-^ colonies was computed, per plate, for each strain/cassette combination. To assess genomic DNA (gDNA) removal, four individual GFP^-^ colonies (three from the 90kbp locus) were picked per strain. gDNA was extracted and prepared for whole genome sequencing as described below (*Sample and library preparation for whole genome sequencing* and *WGS data analysis*).

The progenitor strain for SLIM-mediated genome reduction in *E. coli* (R_0_) was generated from these experiments; coming from the set of colonies picked following Cas3-mediated DNA removal at the 90kbp non-essential span locus. Colony 1 was chosen to be the progenitor strain after verifying the removal of the endogenous chromosomal occurrence of the 34-bp Cas3 target sequence utilized in both the original and modified target cassettes (R_0_, **Extended Data Fig. 1b**).

### SLIM workflows

Genome reduction was performed in parallel across 10 independent lineages over 21 iterative rounds. Each round consisted of (i) pTn5-target cassette delivery and transposition, (ii) screening for target cassette transposition/delivery vector loss, and (iii) Cas3-mediated DNA removal. To initiate Round 1, 20 colonies were selected post-transposition and paired to establish 10 lineages. In subsequent rounds, two colonies per lineage were harvested from the preceding round’s terminal plates (or glycerol stocks) to serve as the template for the next iteration. *P. putida* was cultured at 30°C for all steps.

pTn5 delivery strategies:

- **Phage transduction (1 day/2 days):** Phage transduction was performed using an engineered P1 bacteriophage/phagemid system^51^. Phage particle preparations were performed as previously described^51^. As described by Hur *et al*^48^, phagemid designs were modified to replicate via the π-replicator (piR)/R6Kɣ system, to be unable to replicate in cells targeted for minimization. Target cells were pre-induced in selective ePLM (gentamicin) supplemented with 0.5% arabinose (4h, 37°C) and infected with pTn5-v11 phage lysate for 40 min (20 min shaking, 20 min static). Reactions were quenched with SOC containing 200 mM sodium citrate, recovered in selective LB (gentamicin, hygromycin) + 0.5% arabinose for 3h, and plated. The alternative, 2-day, protocol utilized overnight pre-induction in selective LB before shifting to ePLM for infection.
- **Conjugation (2 days):** Conjugation was mediated by an incP conjugation system derived from pRK24^52^. Overnight cultures of recipient cells (gentamicin, pre-induced with 0.5% arabinose) and donor *E. coli* piR_genRK24 + pTn5-v5 (carbenicillin + chloramphenicol + kanamycin + hygromycin + 2% glucose) were harvested and combined at a 1:5 ratio (recipient:donor). Conjugation was performed on LB agar spots at 30°C for 2h. Cells were recovered in selective LB (gentamicin, hygromycin) for 2h and plated on selective LB agar containing 0.5% arabinose for transposition induction.
- **Electroporation (3 days):** Target cells were grown overnight in 3 mL selective LB (gentamicin). 150 uL of each overnight culture was combined and used to inoculate a new culture in selective LB which was to grown to OD_600_ 0.45–0.6, washed thrice in ice-cold H_2_O, and transformed with 3.5 µL pTn5-v2 plasmid (2.5 kV, 2 mm cuvette). After 1h recovery in LB + 0.5% glucose, cells were grown overnight in selective LB (gentamicin, hygromycin). The following day, cultures were back diluted to OD_600_ 0.05 and transposition was induced with 0.5% arabinose (2h), followed by *sacB*-mediated counterselection against the pTn5-v2 backbone using 7.5% sucrose (4h, 30°C) before plating on selective LB agar (gentamicin, hygromycin) plates.

Universal DNA removal:

- **Cas3-mediated DNA removal (2 days):** Following transposition, four GFP^+^ colonies per lineage were resuspended and spotted on selective LB agar to screen for pTn5 loss. Simultaneously, colonies were independently expanded overnight in selective LB. For Cas3 induction, 1.5 mL of each culture from two verified colonies were pooled, harvested, and resuspended in 3 mL selective LB (gentamicin) containing 0.3% rhamnose. Following 5.5h incubation, cells were plated on selective LB agar containing 5 mM 4-chlorophenylalanine to select for cassette loss. For *P. putida*, selection utilized gentamicin-only plates, with GFP^-^ colonies identifying successful deletion events.

### Target cassette landing site identification

Delivery and transposition assays were coupled with a modified transposon sequencing (TnSeq) protocol^53^ to identify genomic landing sites for the target cassette. Delivery, using phage transduction, conjugation or electroporation, and transposition assays were performed as described above. Approximately 200 colonies were selected from post-transposition plates, per delivery method, pooled for analysis. gDNA was extracted according to the manufacturer’s instructions (QIAGEN™ DNeasy Blood & Tissue Kit cat # 69504). Concentrations of gDNA were quantified with the Invitrogen Qubit™ 4 Fluorometer. DNA libraries were prepared for modified TnSeq as previously described^53^. Pooled library denaturation and flow cell loading was performed per manufacturer instructions. Prior to sequencing, a custom sequencing primer was spiked into the reagent kit at a final concentration of 0.5 µM to enable targeted sequencing of enriched transposon sequences within the libraries. Sequencing was performed on an Illumina MiSeq using MiSeq reagent kit v3 (600-cycles) with 300 bp paired-end reads and automated demultiplexing and adapter trimming.

Output reads were first aligned to a target cassette reference BWA^54^. All reads which aligned to the target cassette reference were collected using reformat.sh^55^ and then aligned to a reference *E. coli* DH10B genome (retrieved from NCBI RefSeq, assembly GCF_000019425.1), once again using BWA. Locations of aligned reads were annotated then plotted using the pyCirclize python package^56^.

### Sample and library preparation for whole genome sequencing (WGS)

WGS of all strains was performed as previously described^53^. Briefly, bacterial cultures were grown in LB media supplemented with the appropriate antibiotics until confluent. Cultures were harvested and gDNA was extracted from the pellets using DNeasy Blood & Tissue Kit (QIAGEN™ cat # 69504) per manufacturer instructions. Concentrations of gDNA were quantified with the Invitrogen Qubit™ 4 Fluorometer using the 1x dsDNA High Sensitivity (HS) assay kit (Thermo Fisher Scientific cat # Q33231). For library preparation, Nextera™ DNA Flex Library Prep Kit (Illumina) was used per manufacturer instructions to tag, barcode, amplify and add index primers (i5 and i7) to the library. 200 ng of starting gDNA was used per barcoded library. Prepared libraries were quantified with the Qubit™ 4 Fluorometer before pooling 15 ng per barcoded library. Pooled library denaturation and flow cell loading was performed per manufacturer instructions. Sequencing was performed on an Illumina MiSeq using MiSeq reagent kit v3 (600-cycles) with 300 bp paired-end reads and automated demultiplexing and adapter trimming.

### WGS data analysis

Sequencing reads from whole-genome samples were aligned to expected reference genomes using the BWA short-read sequence aligner^54^. Alignment files were further processed with Samtools^57^ and deepTools^58^ was used to estimate RPGC throughout the expected reference genome in 10-kbp bins. Regions of low to zero coverage were analyzed in higher resolution using the Integrative Genomics Viewer (IGV)^59^. Exact endpoints of removed DNA segments were identified using IGV, to single nucleotide resolution.

### Coverage plots

Annotated regions of removed DNA, for each genome-reduced strain, were used to generate coverage plots. Each nucleotide, per strain, was assigned a 0 if removed or a 1 retained. Resulting values were plotted for each genome-reduced strain independently and as the average across all genome-reduced strains.

### Deletion tolerance assessment, by function

To assess deletion tolerance based on function we grouped genes into functional groups based on COG group designation. The percentage of genes deleted in each group we computed on a per strain basis. For the initial assessment, qualitative comparisons of the percentage of deleted genes across genome-reduced strains per group were performed. The median percentage of deleted genes was computed for each group.

### Deletion tolerance assessment, by genomic location

Prior to conducting deletion simulations, deletion probabilities were generated for individual genes by first computing the average coverage per nucleotide based on deletions generated in the 10-member library through 21 rounds of SLIM. Round 0 deletions and deletions in one half of the tandem repeat section were excluded from these analyses. For any nucleotide in the genome, if the nucleotide has not been deleted for a given genome-reduced strain, it was given a value of 1. If the nucleotide has been deleted, it was given a value of 0. The average value for each nucleotide is then computed across all genome-reduced strains. Next, the genome was segmented into bins based on the desired final number of bins (1, 10, 100 and 500). The average coverage per bin was computed and converted to average deletion probability by subtracting average coverage from 1. Each gene in a particular bin was then assigned a deletion probability equivalent to the average deletion probability of that bin.

Using computed deletion probabilities genome-wide deletions were simulated on a per gene basis. For each gene a random number between 0 and 1 was generated. If the randomly generated number was less than the deletion probability, that gene was indicated as deleted in the simulation. Otherwise, the gene was indicated as retained. Deletion predictions were accumulated across 10 simulations for each prediction assessment and the result of each simulation was recorded independently per gene for each simulation. For randomized deletion probability predictions, computed deletion probabilities were scrambled and randomly assigned to individual genes. Randomized deletion probability predictions were carried out in the same manner as described for computed deletion probability predictions.

Each gene was then randomly assigned to one of 25 groups. The percentage of deleted genes was computed, per group, based on observed deletions for each genome-reduced strain and based on simulation outputs utilizing each of computed and randomized deletion probability predictions. For each group the set of the percentage of deleted genes based on observed deletions was compared to each of the percentage of deleted genes sets from simulated deletions based on computed deletion probability predictions and randomized deletion probability predictions. The coefficient of determination (R^2^) was computed using the r2_score function from scikit-learn v1.3.0^60^, for the observed set and each predicted set independently, on a per-group basis.

### Sample preparation for transcriptomic analysis

*E. coli* were grown overnight in 10 mL LB + gentamicin to saturation. The next day, 1 mL of overnight culture was collected, harvested and resuspended in 100 uL of ice cold TE buffer. 2 uL Readylyse lysozyme (LGC Biosearch Technologies, #R1810M) was added to each sample. Samples were then incubated on a shaking heat block for 10 minutes at 25°C and 500 RPM. Following incubation each sample was mixed with 300 uL lysis buffer and RNA extractions were performed according to manufacturer’s instructions (NEB Monarch^(R)^ Total RNA Miniprep Kit, #T2010S). Extracted RNA was quantified using Invitrogen Qubit™ 4 Fluorometer using the RNA High Sensitivity (HS) assay kit (Thermo Fisher Scientific cat # Q32852). rRNA depletion was performed and sequencing libraries from extracted RNA samples were prepped and sequenced on an Illumina NextSeq2000 with 50 bp paired-end reads by the Millard and Muriel Jacobs Genetics and Genomics Laboratory at the California Institute of Technology. RNA-Seq experiments were performed with three biological replicates per strain.

### Sample preparation for proteomic analysis

Three biological replicates per strain were grown overnight in 10 mL LB + gentamicin to saturation. Overnight cultures were harvested and resuspended in 5% sodium dodecyl sulfate (Sigma-aldrich) in 50 mM HEPES, and were homogenized using BeatBox (Preomics) for 10 min under ‘High’ settings. Protein concentration was measured using Pierce BCA protein assay kit (Pierce), and 100 μg of protein was used for further sample preparation. The samples were reduced using 5 mM tris(2-carboxyethyl)phosphine (MilliporeSigma) under 55°C for 10min, and then alkylated with chloroacetamide (MilliporeSigma) under room temperature for 15min. The samples were further acidified to a final concentration of 2.5% phosphoric acid, and 25 μl of sample was combined with 165 μl of 90% methanol with 10% of 1 M triethylammonium bicarbonate (TEAB, Thermo Scientific) according to previous protocol^61^. The samples were then loaded onto S-traps (Protifi), and the S-traps were washed using the same buffer for loading (90% methanol with 10% of 1 M TEAB) for 3 times. For each loading or washing step, the S-traps were centrifuged at 4000 g for 30 seconds to remove the elute. After washing steps, 20 μl of 100 mM TEAB containing 10 μg of TPCK-trypsin (Thermo Scientific) was added into each sample, and the digestion was allowed overnight. The digested peptides were eluted using 40 μl of 50 mM TEAB in water, 0.2% formic acid in water, and 50% acetonitrile in water sequentially with 4000 g centrifugation for 1 min for each elution. The elutes for 3 steps were pooled together and dried using a refrigerated CentiVap concentrator (Labconco). The dried samples were stored in -20°C before resuspended in mobile phase A (2% acetonitrile, 0.2% formic acid, and 97.8% water) for LC-MS/MS analysis.

### LC-MS/MS for proteomic analysis

For proteomic samples, LC-MS/MS experiments were performed by loading 500 ng sample onto an EASY-nLC 1200 (ThermoFisher Scientific, San Jose, CA) connected to an Q Exactive HF Quadrapole – Orbitrap Hybrid mass spectrometer (Thermo Fisher Scientific, San Jose, CA). Peptides were separated on an Aurora Ultimate XT UHPLC column (25 cm × 75 μm, 1.6 μm C18, AUR3-25075C18-XT, Ion Opticks) with a flow rate of 0.35 μL/min and for a total duration of 131 min. The gradient was composed of 3% Solvent B for 1 min, 3–19% B for 72 min, 19–29% B for 28 min, 29–41% B for 20 min, 41–95% B for 3 min, and 95–98% B for 7 min. Solvent A consists of 97.8% H2O, 2% ACN, and 0.2% formic acid, and solvent B of 19.8% H2O, 80% ACN, and 0.2% formic acid. MS1 scans were acquired with a range of 375–1500 Th in the Orbitrap at 60 k resolution. The maximum injection time was 15 ms, and the AGC target was 3 × 106. MS2 scans were acquired at 30 k resolution with a first scan mass as 100 Da. The maximum injection time was 45 ms, and the AGC target was 3 × 106. The isolation window was 1.2 Th, collision energy was 28 NCE, and loop count was 12. Other global settings were set to the following: ion source type, NSI; spray voltage, 2000 V; ion transfer tube temperature, 300°C. Method modification and data collection were performed using Xcalibur software (Thermo Scientific).

### Transcriptomic and proteomics data analysis

RNA sequencing output reads were aligned to a reference transcriptome (retrieved from NCBI RefSeq, assembly GCF_000019425.1) and transcript abundances were quantified using Kallisto^62^; abundances, in the form of transcripts per million (TPM), were output for further analysis.

Proteomic analysis was performed using Proteome Discoverer 2.5 (PD 2.5, Thermo Scientific) software, and SequestHT with Percolator validation. The raw data was searched against *Escherichia coli* proteome (retrieved from NCBI RefSeq, assembly GCF_000019425.1). Percolator FDRs were set at 0.001 (strict) and 0.005 (relaxed). Peptide FDRs were set at 0.001 (strict) and 0.005 (relaxed), with medium confidence and a minimum peptide length of 6. Carbamidomethyl (C) was set as a static modification; oxidation (M) was set as a dynamic modification; acetyl (protein N-term), Met-loss (Protein N-term M) and Met-loss + acetyl (Protein N-term M) were set as dynamic N-Terminal modifications.

Differential expression of transcriptomic and proteomic data was assessed using in-house Python scripts. Abundances were median-normalized and T-statistics/p-values computed via SciPy v1.16.0^63^. Thresholds: p-value < 0.05 and |fold-change| ≥ 1. Pathway lists (KEGG, amiGO, stringdb)^35,36,64^ were scored as the sum of log_2_FC values, with deleted genes either assigned a deletion penalty of –10 or removed from the assessment, as indicated. PCA utilized scikit-learn v1.3.0^60^.

### Transcript and protein expression analyses

Following normalization, transcript counts and corresponding protein abundance measurements from each strain were averaged across biological replicates, independently, on a per strain basis. COG group sums were normalized by COG group size. Individual COG group ratios were computed by dividing the group totals for transcripts or proteins by the total transcripts or proteins counted across all groups for each individual strain.

### Fitness assessments

Fitness assessments were conducted in 4 media; standard media (Luria-Bertani, Miller’s LB Broth Base, Invitrogen, #12795-084), rich media (Terrific broth, TB, Ag Scientific Inc, #T-2705-500GM), minimal media (M9 mineral medium, MM)^38^, and standard media supplemented with 2% glucose (LBg). For fitness assessments conducted in all media, the genome reduction starting strain *E. coli* DH10B (pCas3-, Ec_ss_), the genome-reduced lineage library progenitor (R_0_), and derivatives of all ten genome-reduced lineages, each containing pCas3, were grown overnight from glycerol stocks in 3 mL LB + gentamicin (streptomycin for Ec_ss_). The next day, overnight cultures were pelleted and washed (2x) with the appropriate medium. 220 uL media + gentamicin (streptomycin for Ec_ss_) was then added to individual wells in a round-bottom 96 well plate. Each well was then inoculated with one of the twelve overnight cultures to a density of OD_600_ 0.05, three wells per strain. Media was added to two additional wells to serve as blanks in downstream analyses. The plate was incubated in the plate reader (Tecan Spark® Multimode Microplate Reader) for 24 hours at a constant temperature of 37°C. OD_600_ measurements were taken every 15 minutes with shaking incubation implemented in between measurements with an amplitude of 2.5 mm and a double-orbital path.

### Data reporting

No statistical methods were used to predetermine sample size. The experiments were not randomized, and investigators were not blinded during experiments and outcome assessment.

## Data availability

The DNA and RNA sequencing data generated in this study have been deposited in the NCBI BioProject database under BioProject number PRJNA1505730. MS data generated in this study have been deposited in the MassIVE database and can be accessed through the reviewer account: MSV000102858_reviewer with password: 7e7zp6e6u67HEors. Scripts used for data analysis are available upon request.

## Acknowledgements

We thank Prof. Shelley M. Payne from the University of Texas at Austin for generously providing Shigella flexneri CFS100. This work was supported by the Millard and Muriel Jacobs Genetics and Genomics Laboratory at California Institute of Technology. The authors would like to thank Dr. Charles Sanfiorenzo for insight and advice into many aspects of this study, Dr. Russell Swift and Dr. Jianyi Huang for insight into the early stages of development of SLIM, Aline Milach Teixeira for assistance with experiments, Dr. Zach Martinez for thoughtful and thorough review of the manuscript, and Nikki Ma for unwavering support.

## Author contributions

M.L. and K.W. conceived this work and designed experiments. B.Q. designed LC/MS experiments. M.L., B.Q., I.M., G.A., and M.B. conducted experiments and collected data. M.L. and B.Q. generated computational scripts used for DNA, RNA and protein sequencing data analysis and interpreted the results of this work. M.L. and K.W. wrote the manuscript and B.Q., I.M., G.A., M.B., and T.F.C. participated in discussion and provided feedback.

---

**Extended Data Figure 1.**
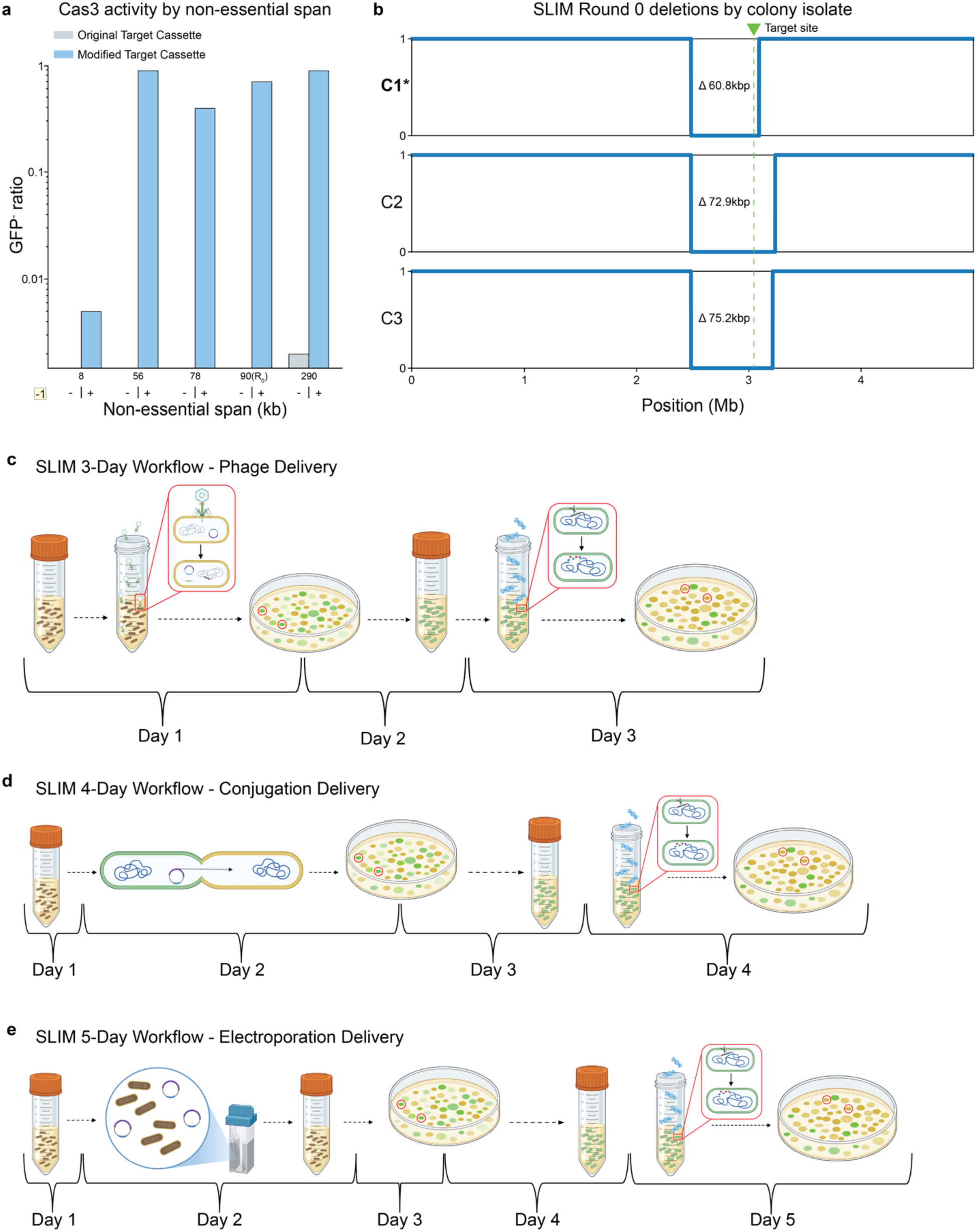
(a) Bar plots displaying the ratio of GFP^-^ colonies counted on plates following Cas3 induction in strains containing either the original (gray) or modified (light blue) target cassette integrated into one of 5 loci, per strain, indicated on the x-axis by non-essential span. Utilization of pheS^mut^ and 4-chlorophenylalanine for counterselection (-1, +) against maintenance of the target cassette is indicated on the x-axis. (b) Coverage plot depicting deletions generated in three independent colonies during “round 0” of SLIM. The progenitor strain from which the library of genome reduced *E. coli* was derived was generated during this round (**C1\***). (c-e) Workflow schematics depicting the 3-day (phage transduction), 4-day (conjugation), and 5-day (electroporation) SLIM protocols. Detailed explanations of each workflow can be found in the Methods section (SLIM workflows).

**Extended Data Figure 2.**
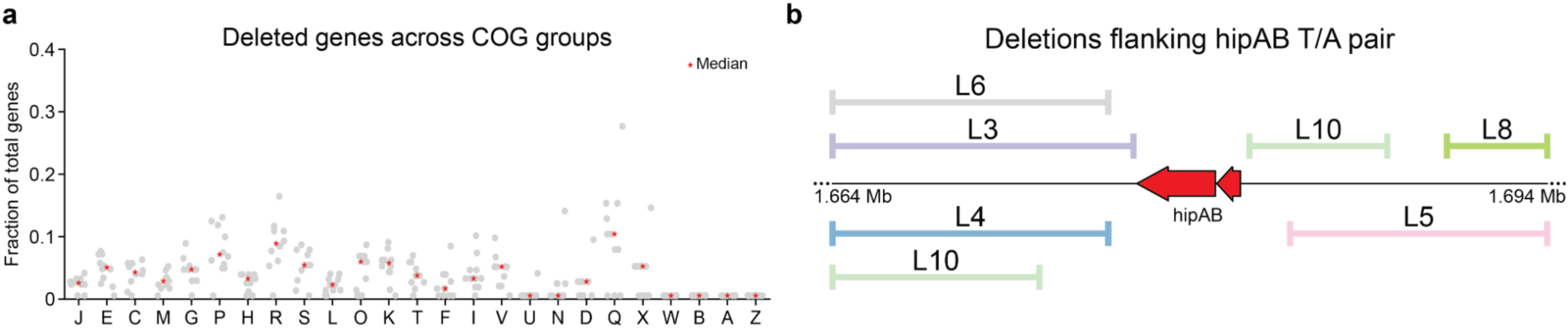
(a) Scatter plot displaying the fraction of deleted genes per derivative of genome-reduced lineages (gray) and the median fraction of deleted genes (*, red) for each COG group. COG groups Key: [A] RNA processing and modification; [B] Chromatin structure and dynamics; [C] Energy production and conversion; [D] Cell cycle control, cell division, chromosome partitioning; [E] Amino acid transport and metabolism; [F] Nucleotide acid transport and metabolism; [G] Carbohydrate transport and metabolism; [H] Coenzyme transport and metabolism; [I] Lipid transport and metabolism; [J] Translation, ribosomal structure and biogenesis; [K] Transcription; [L] Replication, recombination and repair; [M] Cell wall/membrane/envelope biogenesis; [N] Cell motility; [O] Posttranslational modification, protein turnover, chaperones; [P] Inorganic ion transport and metabolism; [Q] Secondary metabolites biosynthesis, transport and catabolism; [R] General function prediction only; [S] Function unknown; [T] Signal transduction mechanisms; [U] Intracellular trafficking, secretion, and vesicular transport; [V] Defense mechanisms; [W] Extracellular structures; [X] Mobilome: prophages, transposons; [Z] Cytoskeleton. (b) Genome schematic indicating the deletions that were generated using SLIM, immediately flanking the toxin-antitoxin pair hipAB.

**Extended Data Figure 3.**
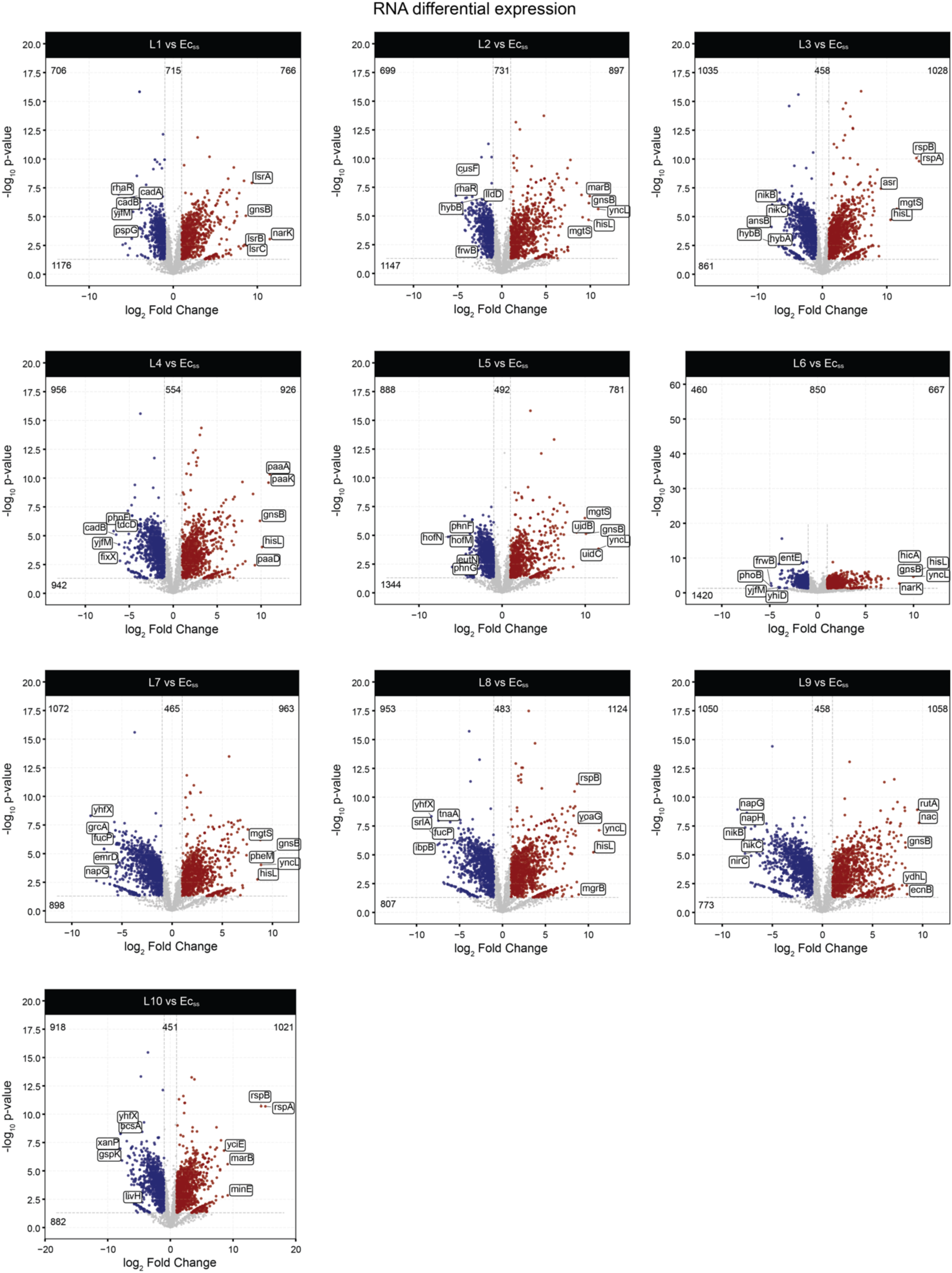
Volcano plots displaying the differential expression of every transcript identified and counted, per derivative of genome-reduced lineages, as compared to Ec_ss_. Upregulated transcripts are indicated in red, downregulated transcripts are indicated in blue, and insignificant transcripts are indicated in gray.

**Extended Data Figure 4.**
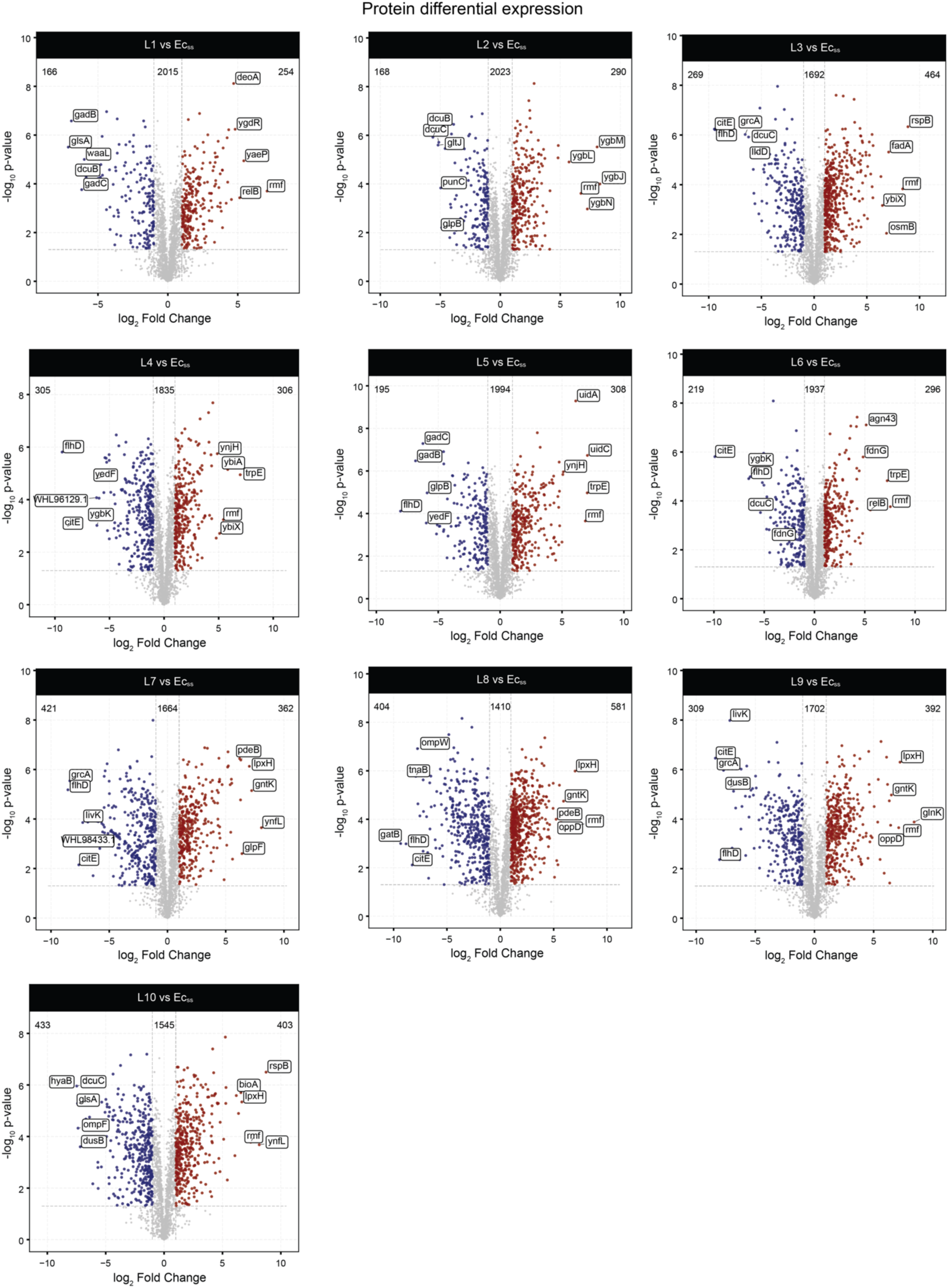
Volcano plots displaying the differential expression of every protein identified and counted, per derivative of genome-reduced lineages, as compared to Ec_ss_. Upregulated proteins are indicated in red, downregulated proteins are indicated in blue, and insignificant proteins are indicated in gray.

**Extended Data Figure 5.**
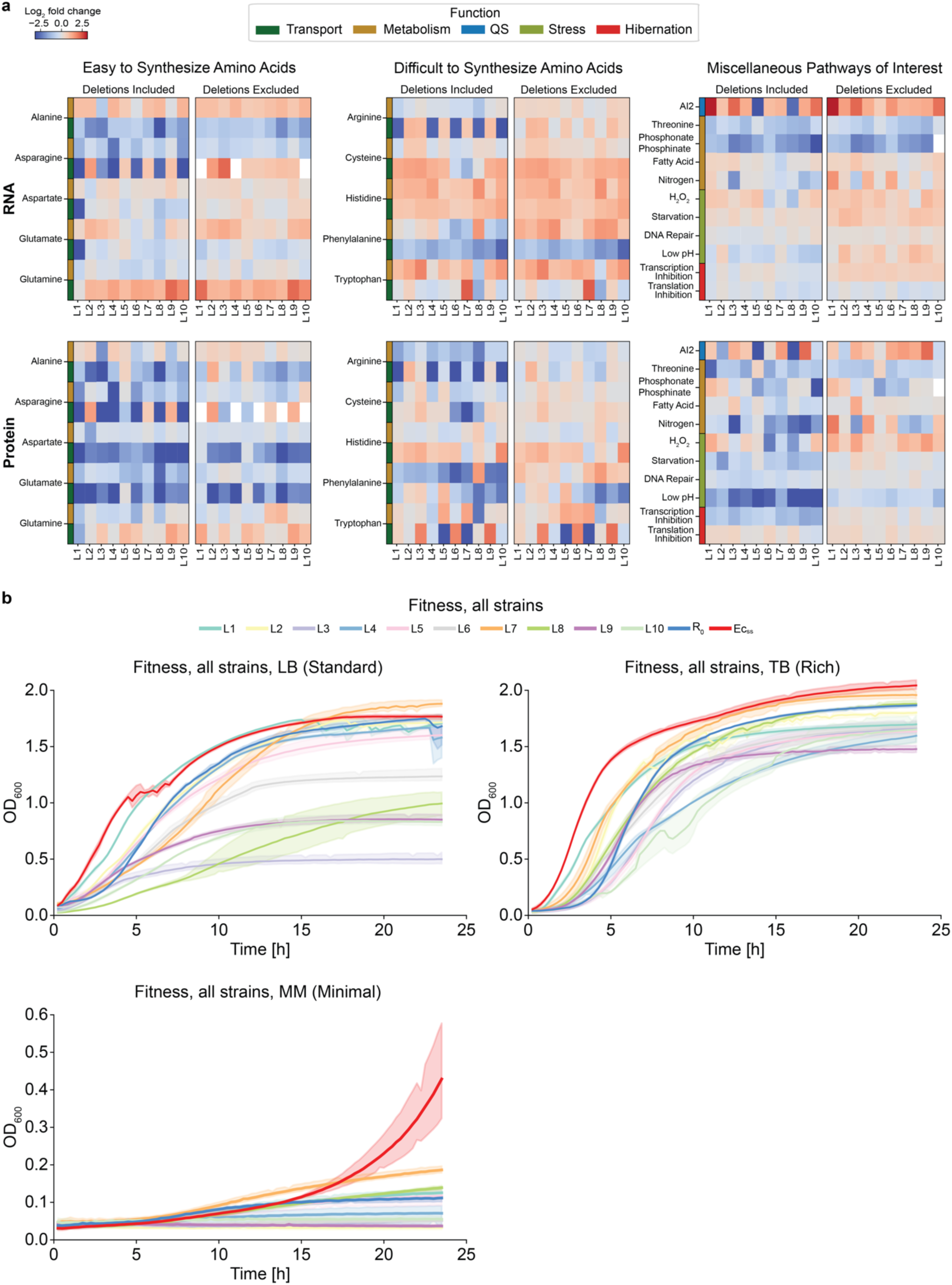
(a) The differential expression of select pathways, expressed as total pathway expression, for each derivative of genome-reduced lineages as compared to Ec_ss_, according to either transcriptomic (top row) or proteomic (bottom row) sequencing output. Heatmaps are broken up into three categories: easy to synthesize amino acids (1-2 steps from a central metabolism intermediate), difficult to synthesize amino acids (4+ steps) and miscellaneous pathways of interest. Each category is further divided into two subcategories for which parallel analyses were conducted: ‘deletion penalty’ and ‘no deletion penalty.’ ‘Deletion penalty’ computations apply the lowest observed log_2_ fold change value to deleted genes in total pathway expression computations. ‘No deletion penalty’ computations simply exclude deleted genes from total pathway expression computations. General function information is indicated by colored rectangles. (b) Fitness curves for derivatives of all genome-reduced *E. coli* lineages, the progenitor strain (R_0_), and the starting strain (Ec_ss_) in three media spanning a gradient of nutrient capacity: standard (LB), rich (TB), and minimal (MM).

**Extended Data Figure 6.**
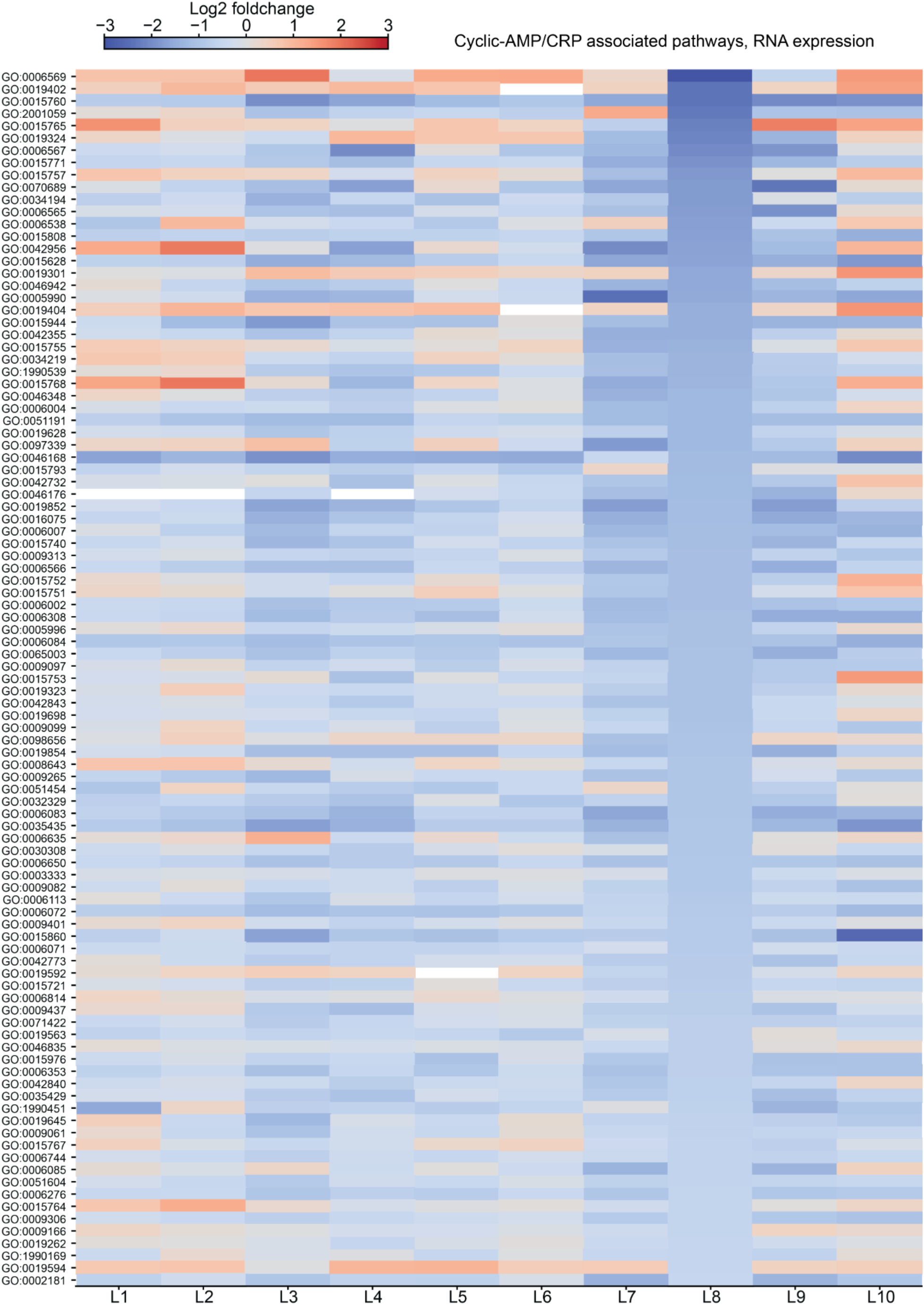
Differential expression, derived from transcriptomic data, of pathways which contain genes that are regulated by cyclic-AMP-CRP, across all genome-reduced *E. coli* lineages. The top 100 most downregulated pathways based on transcriptomic data for L8 are displayed. Pathways are indicated by gene ontology (GO) terms.

**Extended Data Figure 7.**
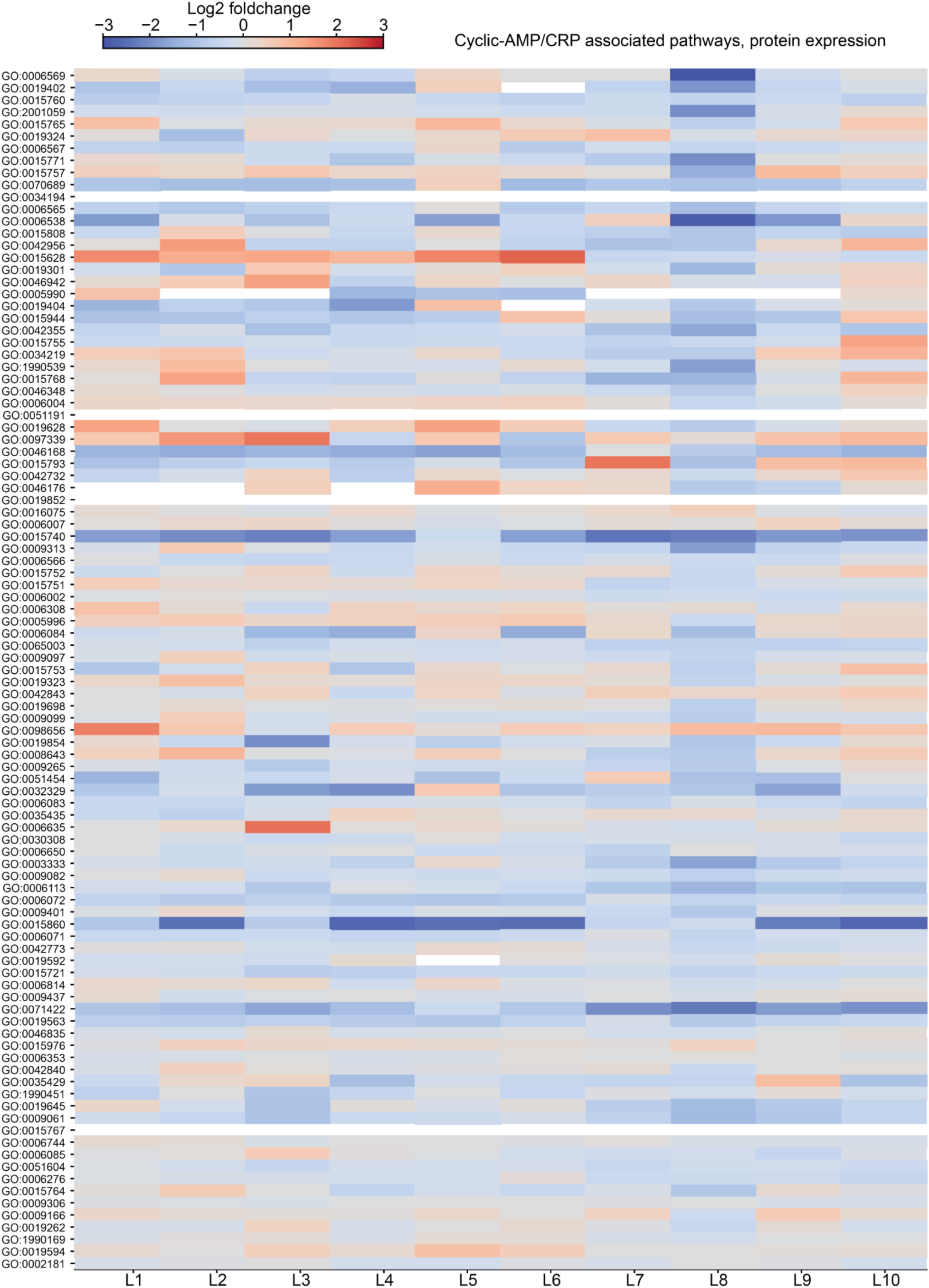
Differential expression, derived from proteomic data, of pathways which contain genes that are regulated by cyclic-AMP-CRP, across all genome-reduced *E. coli* lineages. The top 100 most downregulated pathways based on transcriptomic data for L8 are displayed. Pathways are indicated by gene ontology (GO) terms.

**Extended Data Figure 8.**
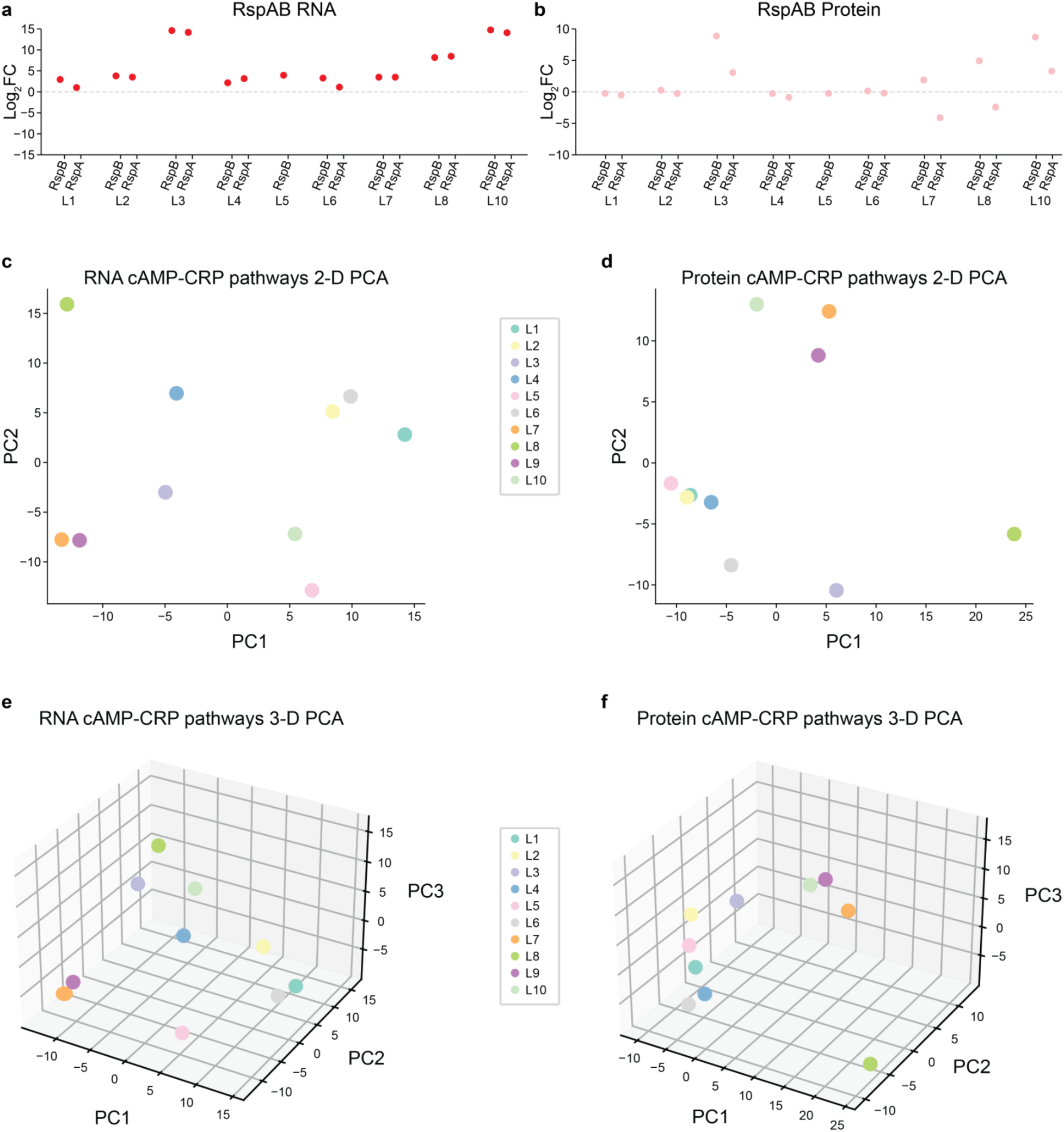
(a) RspAB differential RNA expression for each derivative of genome-reduced lineages, as compared to Ec_ss_. (b) RspAB differential protein expression for each derivative, as compared to Ec_ss_. (c) 2-D principle component analysis (PCA) plot depicting the values of the top two principle components for each derivative of genome-reduced *E. coli* lineages, determined by PCA of transcriptomics-derived cyclic-AMP-CRP associated pathway scores. (d) 2-D principle component analysis (PCA) plot depicting the values of the top two principle components for each derivative of genome-reduced *E. coli* lineages, determined by PCA of proteomics-derived cyclic-AMP-CRP associated pathway scores. (e) 3-D principle component analysis (PCA) plot depicting the values of the top three principle components for each derivative of genome-reduced *E. coli* lineages, determined by PCA of transcriptomics-derived cyclic-AMP-CRP associated pathway scores. (f) 3-D principle component analysis (PCA) plot depicting the values of the top three principle components for each derivative of genome-reduced *E. coli* lineages, determined by PCA of proteomics-derived cyclic-AMP-CRP associated pathway scores.

**Extended Data Table 1:** Coordinates and sizes of all deletions generated using SLIM in this study.

| Strain | Coordinates | Size |
| --- | --- | --- |
| All Strains (rd0) | 248,478..309,298 | 60,821 |
| L1 | 514,313..627,568 | 113,256 |
| L1 | 735,072..739,721 | 4650 |
| L1 | 910,203..970,768 | 60,566 |
| L1 | 1,446,335..1,472,745 | 26,411 |
| L1 | 1,478,719..1,484,546 | 5828 |
| L1 | 1,522,966..1,531,578 | 8613 |
| L1 | 1,532,070..1,533,576 | 1507 |
| L1 | 1,577,553..1,622,608 | 45,056 |
| L1 | 3,095,393..3,109,178 | 13,786 |
| L1 | 3,202,149..3,214,357 | 12,209 |
| L1 | 3,355,526..3,360,477 | 4952 |
| L1 | 3,589,174..3,607,682 | 18,509 |
| L1 | 3,755,464..3,770,393 | 14,930 |
| L1 | 3,838,007..3,874,032 | 36,026 |
| L1 | 4,168,448..4,186,994 | 18,547 |
| L1 | 4,389,294..4,405,915 | 16,622 |
| L1 | 4,591,361..4,644,364 | 53,004 |
| L2 | 330,543..361,830 | 31,288 |
| L2 | 400,888..402,245 | 1358 |
| L2 | 514,313..627,568 | 113,256 |
| L2 | 832,845..845,530 | 12,686 |
| L2 | 855,172..863,066 | 7895 |
| L2 | 1,348,860..1,397,857 | 48,998 |
| L2 | 1,589,819..1,599,674 | 9856 |
| L2 | 1,755,492..1,772,031 | 16,540 |
| L2 | 1,830,433..1,831,124 | 692 |
| L2 | 2,203,830..2,206,699 | 2870 |
| L2 | 2,557,511..2,567,725 | 10,215 |
| L2 | 2,850,764..2,896,946 | 46,183 |
| L2 | 3,119,399..3,125,312 | 5914 |
| L2 | 3,635,303..3,641,667 | 6365 |
| L2 | 3,706,471..3,712,036 | 5566 |
| L2 | 3,842,140..3,852,356 | 10,217 |
| L2 | 3,855,898..3,874,495 | 18,598 |
| L2 | 4,599,279..4,640,717 | 41,439 |
| L3 | 514,313..627,568 | 113,256 |
| L3 | 1,218,350..1,218,424 | 75 |
| L3 | 1,251,185..1,280,884 | 29,700 |
| L3 | 1,349,013..1,397,832 | 48,820 |
| L3 | 1,445,493..1,448,188 | 2696 |
| L3 | 1,483,658..1,522,728 | 39,071 |
| L3 | 1,540,968..1,678,465 | 137,498 |
| L3 | 1,712,393..1,731,497 | 19,105 |
| L3 | 1,848,997..1,866,880 | 17,884 |
| L3 | 2,889,302..2,897,480 | 8179 |
| L3 | 3,195,373..3,200,817 | 5445 |
| L3 | 3,202,149..3,214,162 | 12,014 |
| L3 | 3,457,967..3,462,671 | 4705 |
| L3 | 3,737,134..3,749,152 | 12,019 |
| L3 | 3,791,428..3,819,137 | 27,710 |
| L3 | 3,855,935..3,875,015 | 19,081 |
| L3 | 4,028,966..4,035,978 | 7013 |
| L3 | 4,601,195..4,619,652 | 18,458 |
| L4 | 331,522..336,987 | 5466 |
| L4 | 514,313..627,568 | 113,256 |
| L4 | 755,983..756,488 | 506 |
| L4 | 786,639..804,587 | 17,949 |
| L4 | 902,435..974,939 | 72,505 |
| L4 | 1,546,851..1,561,198 | 14,348 |
| L4 | 1,590,166..1,622,365 | 32,200 |
| L4 | 1,644,135..1,677,195 | 33,061 |
| L4 | 2,546,095..2,549,761 | 3667 |
| L4 | 2,848,680..2,854,427 | 5748 |
| L4 | 3,192,736..3,200,814 | 8079 |
| L4 | 3,202,149..3,213,275 | 11,127 |
| L4 | 3,585,876..3,604,347 | 18,472 |
| L4 | 3,719,495..3,742,959 | 23,465 |
| L4 | 3,837,936..3,871,180 | 33,245 |
| L4 | 4,159,500..4,167,679 | 8180 |
| L4 | 4,590,283..4,641,938 | 51,656 |
| L5 | 1,682,394..1,698,853 | 16,460 |
| L5 | 1,785,485..1,801,299 | 15,815 |
| L5 | 2,847,422..2,848,646 | 1225 |
| L5 | 3,722,691..3,817,577 | 94,887 |
| L5 | 3,837,759..3,878,532 | 40,774 |
| L5 | 4,157,441..4,167,676 | 10,236 |
| L5 | 4,301,284..4,322,829 | 21,546 |
| L6 | 514,313..627,568 | 113,256 |
| L6 | 755,984..756,499 | 516 |
| L6 | 918,152..964,837 | 46,686 |
| L6 | 1,325,192..1,329,514 | 4323 |
| L6 | 1,508,605..1,551,819 | 43,215 |
| L6 | 1,644,135..1,677,195 | 33,061 |
| L6 | 1,699,220..1,712,420 | 13,201 |
| L6 | 1,758,419..1,764,903 | 6485 |
| L6 | 2,247,288..2,269,694 | 22,407 |
| L6 | 2,546,095..2,549,761 | 3667 |
| L6 | 2,848,680..2,855,825 | 7146 |
| L6 | 3,166,482..3,171,156 | 4675 |
| L6 | 3,214,607..3,241,893 | 27,287 |
| L6 | 3,577,527..3,581,395 | 3869 |
| L6 | 3,585,363..3,603,738 | 18,376 |
| L6 | 3,794,166..3,817,581 | 23,416 |
| L6 | 3,835,502..3,853,420 | 17,919 |
| L6 | 3,995,746..4,002,291 | 6546 |
| L6 | 4,382,247..4,388,525 | 6279 |
| L7 | 447,589..463,418 | 15,830 |
| L7 | 585,548..624,911 | 39,364 |
| L7 | 1,348,736..1,397,676 | 48,941 |
| L7 | 1,414,040..1,416,922 | 2883 |
| L7 | 1,451,456..1,605,432 | 153,977 |
| L7 | 1,906,701..1,907,631 | 931 |
| L7 | 2,082,611..2,107,573 | 24,963 |
| L7 | 2,343,565..2,346,754 | 3190 |
| L7 | 2,412,883..2,418,078 | 5196 |
| L7 | 3,168,874..3,170,543 | 1670 |
| L7 | 3,202,149..3,223,630 | 21,482 |
| L7 | 3,376,480..3,402,553 | 26,074 |
| L7 | 3,647,821..3,657,912 | 10,092 |
| L7 | 3,940,322..3,943,595 | 3274 |
| L7 | 4,028,111..4,034,434 | 6324 |
| L7 | 4,107,426..4,114,268 | 6843 |
| L7 | 4,389,295..4,412,573 | 23,279 |
| L8 | 444,330..475,983 | 31,654 |
| L8 | 514,313..627,568 | 113,256 |
| L8 | 921,252..974,006 | 52,755 |
| L8 | 1,348,644..1,397,893 | 49,250 |
| L8 | 1,689,072..1,729,845 | 40,774 |
| L8 | 1,906,701..1,907,632 | 932 |
| L8 | 2,847,791..2,896,946 | 49,156 |
| L8 | 3,200,768..3,214,415 | 13,648 |
| L8 | 3,572,879..3,588,504 | 15,626 |
| L8 | 3,635,288..3,641,652 | 6365 |
| L8 | 3,642,728..3,658,032 | 15,305 |
| L8 | 3,722,709..3,817,581 | 94,873 |
| L8 | 4,028,111..4,034,434 | 6324 |
| L8 | 4,168,449..4,191,832 | 23,384 |
| L8 | 4,388,776..4,392,174 | 3399 |
| L8 | 4,636,624..4,644,362 | 7739 |
| L9 | 456,607..490,357 | 33,751 |
| L9 | 514,313..627,568 | 113,256 |
| L9 | 647,631..651,623 | 3993 |
| L9 | 669,421..689,982 | 20,562 |
| L9 | 1,324,984..1,330,658 | 5675 |
| L9 | 1,348,769..1,398,180 | 49,412 |
| L9 | 1,447,706..1,503,067 | 55,362 |
| L9 | 1,528,191..1,559,116 | 30,926 |
| L9 | 1,594,464..1,610,270 | 15,807 |
| L9 | 1,726,152..1,750,416 | 24,265 |
| L9 | 2,309,280..2,332,993 | 23,714 |
| L9 | 2,560,066..2,589,699 | 29,634 |
| L9 | 2,755,676..2,783,803 | 28,128 |
| L9 | 3,740,288..3,781,962 | 41,675 |
| L9 | 3,898,535..3,901,288 | 2754 |
| L9 | 4,167,872..4,187,975 | 20,104 |
| L9 | 4,374,147..4,389,130 | 14,984 |
| L9 | 4,637,459..4,644,363 | 6905 |
| L10 | 514,313..627,568 | 113,256 |
| L10 | 911,662..965,084 | 53,423 |
| L10 | 1,503,980..1,674,056 | 170,077 |
| L10 | 1,680,231..1,687,263 | 7033 |
| L10 | 1,713,176..1,730,196 | 17,021 |
| L10 | 1,737,524..1,740,539 | 3016 |
| L10 | 1,780,779..1,791,521 | 10,743 |
| L10 | 1,808,046..1,812,467 | 4422 |
| L10 | 2,557,511..2,567,725 | 10,215 |
| L10 | 3,284,544..3,289,336 | 4793 |
| L10 | 3,457,879..3,462,671 | 4793 |
| L10 | 3,632,495..3,642,271 | 9777 |
| L10 | 3,731,020..3,781,948 | 50,929 |
| L10 | 3,945,724..3,970,479 | 24,756 |
| L10 | 4,156,829..4,167,679 | 10,851 |
| L10 | 4,317,579..4,344,277 | 26,699 |
| L10 | 4,389,294..4,452,419 | 63,126 |
| L10 | 4,637,076..4,641,938 | 4863 |
| SL1 | 1,769,283..1,769,728 | 446 |
| SL1 | 3,105,224..3,144,989 | 39,766 |
| SL1 | 3,478,885..3,479,182 | 298 |
| SL2 | 807,064..859,864 | 52,801 |
| SL2 | 3,121,010..3,152,127 | 31,118 |
| SL2 | 3,965,561..3,967,668 | 2108 |
| SL3 | 500,478..534,804 | 34,327 |
| SL3 | 2,178,448..2,211,256 | 32,809 |
| SL3 | 3,083,001..3,128,297 | 45,297 |
| PL1 | 869,849..880,781 | 10,933 |
| PL1 | 3,820,981..4,068,420 | 247,440 |
| PL1 | 4,790,351..4,824,366 | 34,016 |
| PL2 | 4,531,459..4,619,852 | 88,394 |
| PL3 | 2,285,330..2,293,514 | 8185 |
| PL3 | 3,362,126..3,718,902 | 356,777 |

**Extended Data Table 2:** Deleted essential genes in *E. coli*.

| Strain | Deleted Essential Genes (Any Study<sup>20–22</sup>) | Deleted Essential Genes (Consensus) |
| --- | --- | --- |
| L1 | yagG', 'yncH', 'tdcF' | n/a |
| L2 | yagG' | n/a |
| L3 | yagG', 'yncH', 'ydfO', 'minD', 'ydiL', 'safA',<br>'racR', 'ydaS', 'iraM' | n/a |
| L4 | yncH', 'safA', 'yagG' | n/a |
| L5 | bcsB', 'yagG' | n/a |
| L6 | safA', 'yagG' | n/a |
| L7 | racR', 'ydaS', 'alsK', 'rsmI', 'yagG' | n/a |
| L8 | yagG', 'ydfO', 'bcsB' | n/a |
| L9 | relB', 'yagG', 'glyA', 'dicA' | n/a |
| L10 | yncH', 'yagG', 'ydaS', 'alsK', 'safA', 'ydfO',<br>'racR' | n/a |

